# Bumble bee workers adopt novel behavioral roles and reshape their social networks in the absence of a queen

**DOI:** 10.1101/2025.01.07.630106

**Authors:** Dee M. Ruttenberg, Scott W. Wolf, Andrew E. Webb, Eli S. Wyman, Michelle L. White, Diogo Melo, Ian M. Traniello, Sarah D. Kocher

**Author notes:** Correspondence: Ian M. Traniello < >, Sarah D. Kocher < >. Authors contributed equally to this manuscript (Dee M. Ruttenberg, Scott W. Wolf, Ian M. Traniello, Sarah D. Kocher).

## Abstract

Dominant individuals often structure group organization, but less is known about how social networks reorganize in their absence and how variation among subordinates contributes to collective outcomes. Bumble bees (*Bombus impatiens*) provide an ideal system to study these dynamics: queens typically monopolize reproduction, but in some contexts individual workers can adopt queenlike social roles. Using multi-animal pose tracking, we compared matched queenright and queenless partitions from the same source colonies, quantifying over 80 million social interactions. Queen-less colonies exhibited increased behavioral variation and contained a subset of highly influential workers with elevated movement, spatial centrality, and reproductive activity that was absent in queen-right conditions. The emergence of these individuals coincided with a shift from centralized to decentralized, efficient network architectures. These results demonstrate that queen presence constrains latent worker variation, revealing how individual behavioral differences can scale up to reshape collective social organization in hierarchical societies.

## Introduction

The influence of individuals within a social network can vary dramatically, and some members can play an outsized role in shaping group behavior and organization (Jolles, King, and Killen, 2019; Kralj-Fišer and Schuett, 2014; Cook et al., 2020). Highly influential individuals often serve as hubs in their networks, mediating information flow and maintaining social stability (Caticha, Calsaverini, and Vicente, 2024; Newman, 2018; Freeman, 1977), making their presence particularly consequential for group organization and function (Centola, 2019; McCully and Rose, 2023). However, we know comparatively little about how social systems reorganize in the absence of key individuals, or how latent differences among subordinates contribute to this process.

In social insects, individual variation in behavior and physiology is tightly linked to social context. In eusocial colonies, one or a few individuals monopolize reproduction, while the majority of group members perform supportive tasks as functionally sterile workers. Reproductively dominant queens can suppress worker reproduction through mechanisms ranging from direct physical interactions to chemical signals that inhibit ovarian activation (Van Zweden, 2010; Smith and Liebig, 2017). The queen also plays a central role in structuring group interactions (Gadagkar, Sharma, and Pinter-Wollman, 2022). Yet, in many social insect species, colonies can continue to persist and function for some time following queen loss, raising the question of how social structure is maintained or reorganized in the absence of a central reproductive figure.

Queenless conditions offer a lens into how social context shapes collective behavior. In some species, the absence of a queen is associated with increased behavioral competition and reproductive dominance among workers: some colony members can activate their ovaries and lay eggs, often leading to competition among nestmates (Wilson, 1971; Amsalem, Twele, et al., 2009). These shifts are thought to reflect underlying behavioral and physiological variation among workers that is present across social contexts but typically constrained under queenright conditions (Kocher et al., 2010; Kaatz, Hildebrandt, and Engels, 1992). It remains unclear how such individual differences impact colony social organization or whether the emergence of new behavioral roles reshapes social network structure.

The common eastern bumble bee (*Bombus impatiens*) enables the study of how the presence or absence of a reproductive queen shapes individual behavior and physiology. Moreover, it allows us to examine how this individual-level variation scales up to influence colony-level social organization. Bumble bee colony development begins with a cooperative, eusocial phase characterized by a strong reproductive division of labor between a nest-founding queen and her functionally sterile daughter-workers. Physical contact with the queen is required to prevent workers from becoming reproductively active (Padilla et al., 2016). This contact is thought to, in part, mediate the transmission of chemical cues produced by the queen that signal her reproductive status (Orlova, Treanore, and Amsalem, 2020; Orlova and Amsalem, 2021). The later phase of a colony’s annual life cycle is characterized by a breakdown of reproductive suppression by the queen, and competition emerges between workers to activate their ovaries and lay haploid, male-destined eggs (Goulson, 2010). In *Bombus terrestris*, workers actively monitor queen behavior and brood status and initiate competition when gyne-destined larvae—which develop into the next generation of reproductives—are detected (Alaux, Jaisson, and Hefetz, 2004; Alaux, Jaisson, and Hefetz, 2006). The queen can inhibit gyne development and commit larvae to the worker-destined developmental pathway (Cholé et al., 2022; Franco et al., 2023; Amsalem, Grozinger, et al., 2015), and workers monitor her status to detect when she ceases to exert this influence. Thus, information flow from the queen to the workers is instrumental in the regulation of social structure under queenright conditions. Importantly, the natural transition from eusocial cooperation to competition, can be artificially triggered by the experimental removal of the queen or by separation of workers from a queenright colony (Cnaani, Wong, and Thomson, 2007).

To better understand how queen presence relates to individual behavior and colony-level organization, we used automated multi-animal pose tracking to analyze social interactions in matched queenright and queenless *B. impatiens* partitions. Bumble bee colonies are highly robust to experimental partitioning (Free, 1955; Röseler, 1977; Pandey, Motro, and Bloch, 2020), allowing us to compare groups derived from the same source colony. This matched design enables us to isolate the role of queen presence in shaping group dynamics while controlling for colony-specific and environmental variation.

We implemented a “hybrid” tracking system, NAPS (NAPS is ArUco Plus SLEAP) (Wolf et al., 2023), to capture high-resolution behavioral, spatial, and social interaction data in queenright and queenless contexts. NAPS integrates fiducial marker tracking with pose estimation, enabling high-fidelity tracking of each bee’s identity and their body part positions throughout the duration of an experiment (Pereira et al., 2022; Wolf et al., 2023). Through this, we quantified the spatial dynamics, activity levels, and social interactions of each individual bee over time. We examined how social context shapes variation in worker behavior and physiology, and how these individual-level differences relate to colony-level network structure.

## Results

### Automated Monitoring of Bumble Bee Colonies Unmasks the Dynamics of Physical Contact Interactions

We previously established a hybrid tracking system, NAPS, to automate the monitoring of bumble bee behaviors within a colony setting (Wolf et al., 2023). NAPS integrates pose estimation performed with SLEAP (Pereira et al., 2022) with tag identification using ArUco (Garrido-Jurado et al., 2014). We developed a SLEAP model using training data from five bumble bee colonies, each divided into queenless and queenright partitions and filmed for 96 hours per cohort (Figure 1). Our SLEAP model yielded accurate representations of workers and queens with minimal error (Figure S1, Figure S2). After matching individual instances to their associated ArUco tag, we generated a dataset of each bee’s location. We then filtered our dataset to remove spurious detection events, including bees moving impossibly fast and distances between nodes that were either impossibly large or small (Figure S3). No filtering step was biased between queenless and queenright colonies (Figure S4). We used antennal presence, calculated as the percentage of frames in which all nodes representing the antennae were visible, as a proxy for each bees’ detectability; this value was comparable between queenless and queenright colonies across trials (Figure S5). A single colony (Colony 3, Queenright) had a marginally lower antennal presence compared to other colonies, and we adjusted for this in downstream analyses by normalizing all individual interaction events to antennal presence.

**Figure 1.**
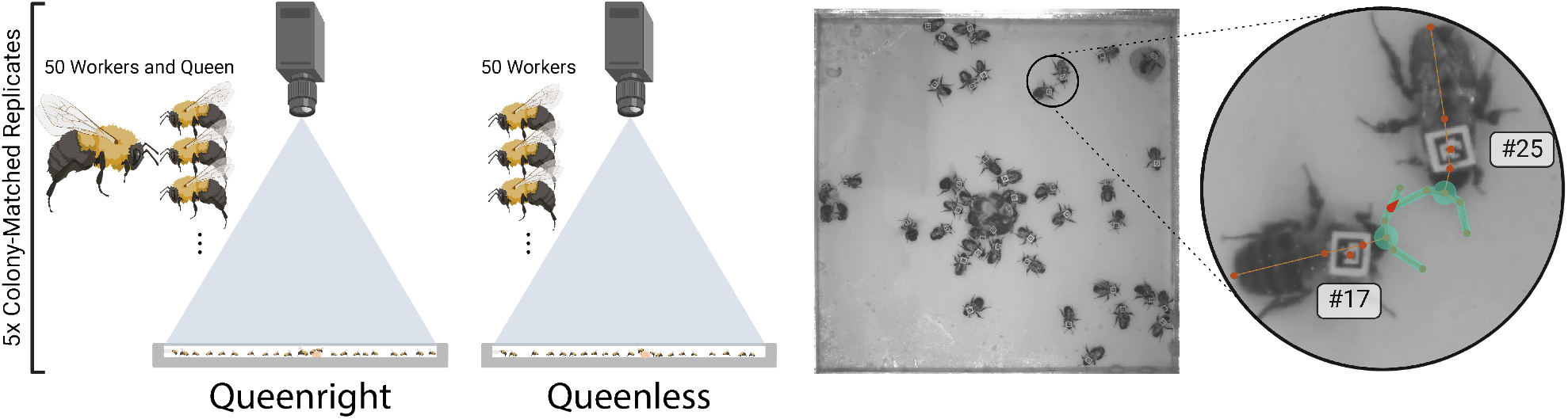
Experimental design and example application of automated behavioral tracking with NAPS. (left) Five colonies of approximately 200 workers and a queen were each split into two partitions: one queenright partition consisting of a queen and 50 (*±*2) workers and another with 50 (*±*2) colony- and size-matched workers. The resulting colonies were imaged under infrared light with controlled temperature (*∼*27 ^*°*^C) and relative humidity (*∼*30%) for four days. (right) Still image from a colony recording: inset shows antennal interaction between adjacent workers, including the localization of nodes and edges (i.e., relevant body parts and their connections, respectively) detected via SLEAP. Unique ArUco identifiers are shown in a white box next to each bee. Antennal overlap, constituting a head-to-head interaction, is shown in red.

One key advantage of pose estimation is its ability to track and quantify different modalities of physical interactions (Traniello and Kocher, 2024). Knowing that bumble bees rely on their antennae to detect physical and chemical cues (Spaethe et al., 2007), we quantified two distinct types of pairwise antennal contact interactions: head-to-head and head-to-body (Figure 1, Figure S6). Despite the bumble bee body being >5x larger than the head, we found head-to-head interactions were significantly more frequent than head-to-body interactions (Figure S6: *t* = 61.21, 81.67, 19.39 for daytime hours (9:00-17:00) in queenright work-ers, queenless workers, and queens respectively; *p <* 2.2*\**10^*−*16^ in all comparisons). This suggests that head-to-head antennal interactions are enriched in bumble bee colonies as a primary means of physical communication, consistent with results from detailed tracking of two-bee pairings (Wang et al., 2022). We detected a slight diel effect in the frequency of head-to-head interactions (Figure 2a, Figure S7), so we restricted our analyses to daytime hours. In total, we quantified over 80 million undirected pairwise interactions across nearly 65 million frames. For over 25 million of these, we could assign a clear initiator based on pre-interaction displacement (see Methods); the remaining interactions had no unambiguous initiator.

**Figure 2.**
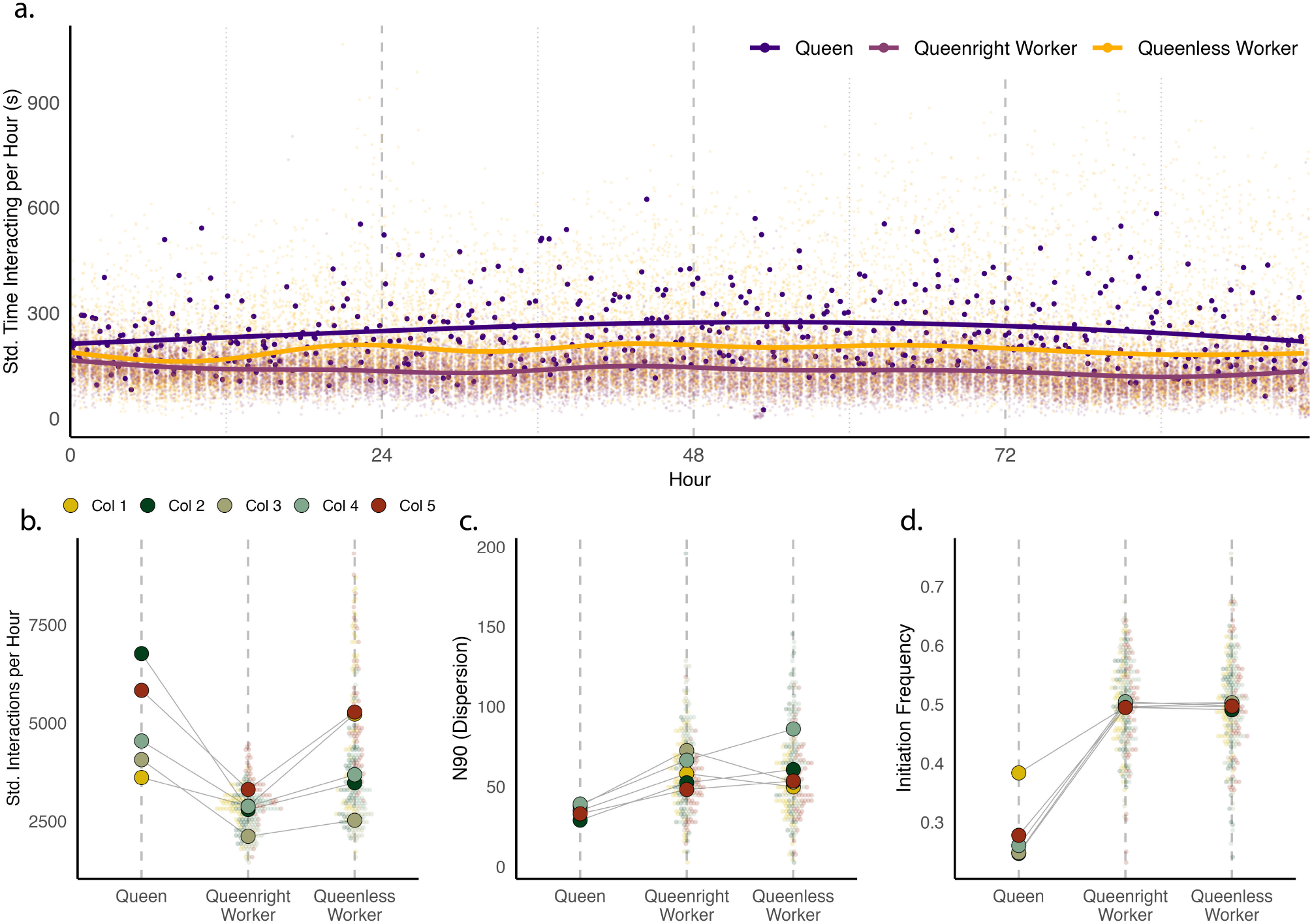
Social interactivity correlates with reproductive state and social condition. (a) Interaction counts (degree centrality) over time aggregated across all colonies within each condition over an hour; counts are standardized to antennal presence for each bee. (b) Standardized interaction counts from (a) averaged for each bee. Queens have significantly higher interaction counts than workers(*χ*^2^ = 26.4, *p* = 2.09 *\** 10^*−*147^). (c) Dispersion metrics as calculated by N90, the smallest number of 200 x 200 pixel squares that represent 90% of a bee’s home range. Queens show significantly lower dispersion than workers (*χ*^2^ = *−*12.8, *p* = 2.39 *\** 10^*−*37^) (d) Proportion of directed interactions in which a given bee is the initiator, as measured by pre-interaction displacement (see Methods). Queens initiate significantly fewer interactions than workers (*χ*^2^ = *−*16.9, *p* = 7.97 *\** 10^*−*63^). For (b-d), large dots represent the mean for a given colony; for queenright and queenless workers, raw data are shown in the background.

### Queens Play a Central Role in the Queenright Social Network

We focused our analyses on head-to-head contact interactions. *B. impatiens* females exhibit an enrichment of head-to-head interactions in our data (Figure S6) and in previously published datasets (Wang et al., 2022; Wolf et al., 2023). These interactions have been historically used to analyze colony behavior (Crall, Gravish, Mountcastle, Kocher, et al., 2018; Wild et al., 2021; Wolf et al., 2023) and are disrupted when bees experience early life social isolation (Wang et al., 2022). We tested the hypothesis that the queen regulates the social environment of the colony (at least in part) by maintaining a high interactivity relative to workers. To do this, we generated weighted, undirected social networks between all bees in the colony for each hour of the experiment; weighting was based on the total time spent in each dyadic interaction. Interaction weights were then standardized to antennal presence for each individual, as described in Methods. This revealed the queen to be the most central bee in the network, interacting much more frequently than nestmate workers, resulting in a higher degree centrality (Figure 2a-b: *χ*^2^ = 26.4, *p* = 2.09 *\**10^*−*147^). This result was also present in unstandardized data (Figure S8a). Relative to interactions between workers, queen-worker interactions were more strongly enriched for head-to-head antennation and longer in duration, suggesting that reproductive status influences physical communication strategies among nestmates (Figure S6, Figure S8b).

Despite the queen interacting and moving more frequently than the workers, she explored relatively less space in the colony (Figure 2c: *χ*^2^ = *−*12.8, *p* = 2.39 *\**10^*−*37^; Figure S8), consis-tent with observations of her primary localization to the brood (Cnaani, Schmid-Hempel, and Schmidt, 2002) and prior work in bumble bees (Jandt, Huang, and Dornhaus, 2009). We further interrogated the dynamics of queen-worker interactivity and found the directionality of these interactions to be imbalanced: queen-worker interactions were initiated by the worker *∼*70% of the time (Figure 2d: *χ*^2^ = *−*16.9, *p* = 7.97 *\**10^*−*63^). Taken in sum, the queen occupies a smaller home range in which she is visited frequently by workers, resulting in a proportionally high interaction rate.

### Individual Behavioral Variation Increases in Queenless Colonies

In light of the queen’s central role in the colony’s social environment, we hypothesized that individual workers would vary more extensively in a queenless setting. To this end, we analyzed queenless partitions of workers collected from the same source colony as the queenright partition. Associations between body size and aggression have been made in the closely related bumble bee species, *Bombus terrestris*, (Princen et al., 2020), a possible confound we addressed by visually size-matching individuals within each cohort and later using marginal cell length, which serves as a robust proxy for body size, to confirm that our visual assessments were accurate (see Methods). Principal component analysis (PCA) performed on interactive (both head-to-head and head-to-body), kinematic, and spatial metrics identified network centrality measures and kinematic variation as the major contributors to the primary axis of variation (i.e., PC1) (Figure 3a, Figure S9). PC1 captured most measures of centrality, including betweenness, closeness, degree, eigenvector centrality, as well as clustering coefficient (Figure S9). Neither of the first two PCs represented variance explained by source colony (Figure S10), suggesting that variance attributed to these PCs was consistently observed across the five colony replicates.

**Figure 3.**
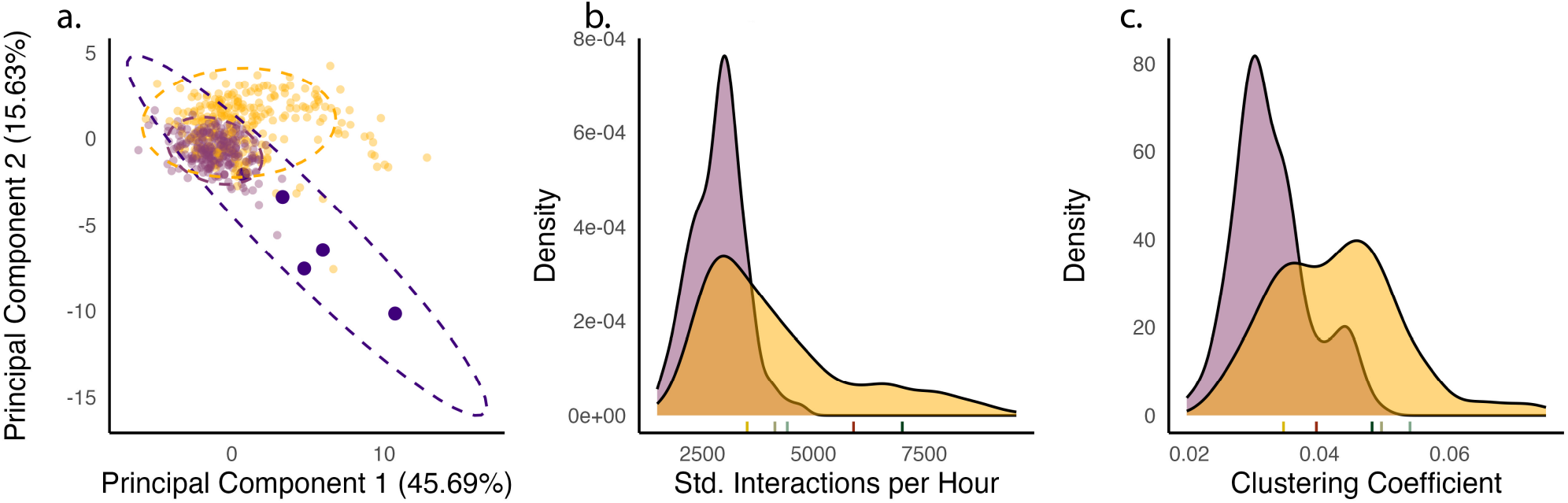
Queenless colonies undergo major shifts in individual- and group-level interaction dynamics. (a) Principal component analysis of behavioral metrics for individual bees in queenright and queenless partitions. Queens and queenright workers are shown in dark and light purple, respectively, and queenless workers are shown in yellow. (b-c) Density plot of (b) standardized interactions per hour (glmmTMB dispersion contrast: *t* = *−*17.92, *p <* 0.0001) and (c) clustering coefficient (glmmTMB dispersion contrast: *t* = *−*15.83, *p <* 0.0001) for queenright and queenless partitions. Queens are indicated in rug plot below each density plot and colored according to source colony. n = 501 workers and five queens from five source colonies.

Using glmmTMB dispersion sub-models (Brooks et al., 2017), we also observed an overall increase in individual variation in queenless colonies. Queenlessness was associated with increased variation in multiple network parameters, including interactivity and tendency to cluster (standardized interaction count: Figure 3b, dispersion contrast: *t* = *−*17.92, *p <* 0.0001; clustering coefficient: Figure 3c, dispersion contrast: *t* = *−*15.83, *p <* 0.0001).

### Queenless Partitions Have Increased Ovarian Development

To understand the biological basis of the observed variation in network metrics, we investigated differences in reproductive physiology between queenright and queenless workers. Using ovary index (Duchateau, 1989; Cnaani, Schmid-Hempel, and Schmidt, 2002), a measure of ovary activation normalized to body size, we found that worker ovary size was on average larger in queenless partitions (Figure S11, Figure S12), consistent with results from other queen removal experiments (Padilla et al., 2016). This suggests a generalized increase in both physiological and behavioral variation among queenless workers compared to queenright workers.

### A Subset of Queenless Workers Express Queen-Like Behavioral and Reproductive Phenotypes

Using the PC1 scores from Figure 3a, which reflect individual variation in interactive, kinematic, and spatial behavior, we identified workers with PC1 values exceeding the highest score observed among queenright individuals from the same source colony. These bees likely expressed the most dramatic shifts in behavior in a queenless context compared to individuals in their paired, queenright partition. This small but highly interactive population of bees contained 37 individuals across all five queenless partitions (two to twelve bees per partition). These individuals were slightly, but significantly, larger than non-outlier nestmates (Figure S13, *χ*^2^ = *−*2.50, *p* = 1.25 *\**10^*−*2^; hub bee worker marginal cell length = 2.684 *±*0.244 mm, non-hub queenless worker mean = 2.642 *±*0.258 mm). Throughout the remainder of the manuscript, we refer to these behaviorally and physiologically distinct individuals as “hub bees,” which reflects their central role in the queenless social network.

Although PC1 is an aggregate of many behavioral metrics (Figure 4a), we independently tested whether hub bees differed from other workers along the individual traits that contributed most to this axis. Hub bees were more interactive than non-hub queenless workers, often having a total number of interactions comparable to or exceeding their source colony’s queen (Figure S14a, *χ*^2^ = 865, *p ≈*0; Figure S9). Similarly to the queen, the hub bees also moved more frequently, were higher in other centrality measures, and were significantly less dispersed than the remaining work-ers (movement percent: *χ*^2^ = 398, *p* = 1.44 *\**10^*−*88^; dispersion Figure S14b: *χ*^2^ = 462, *p* = 1.40 *\**10^*−*102^; betweenness Figure S14c: *χ*^2^ = 398, *p* = 1.78 *\**10^*−*88^). However, other metrics such as interaction initiation rate and movement speed did not differ between hub bees and other workers (Figure 4a), highlighting that the designation of “hub bee” does not simply reflect uniformly elevated activity. Full pairwise comparisons between hub bees and non-hub queenless workers across all behavioral metrics are shown in Figure S14.

**Figure 4.**
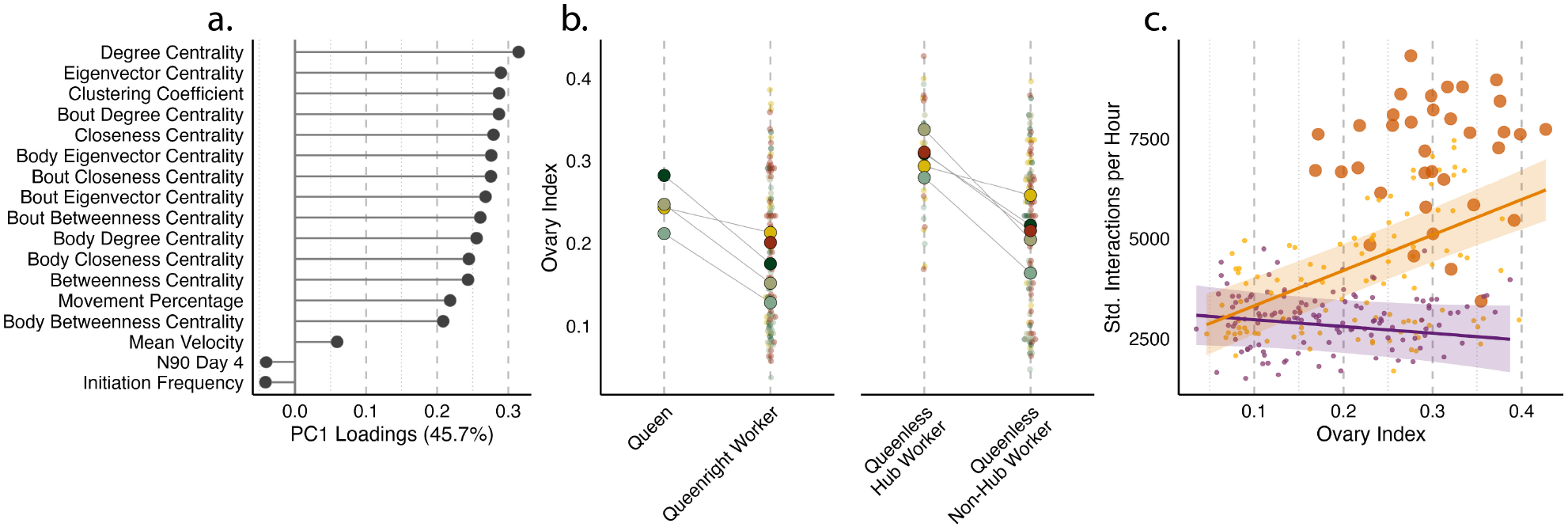
In the absence of a queen, a small subset of workers expresses queen-like behavioral and physiological traits. (a) PC1 loadings for 17 behavioral and spatial parameters. Higher loadings are more associated with “hub bees” (Figure S10), which we defined as workers in a queenless colony with a greater PC1 than all workers in the paired queenright colony. (b) Ovary index (size of the largest ovariole relative to body size) of queens, queenright workers, queenless hub workers, and queenless non-hub workers. Large dots show colony means and smaller dots show individual measurements for ovary index (*χ*^2^ = 25.4, p = 2.88*\** 10^*−*133^). (c) Standardized interactions per hour relative to ovary index in queenright workers (purple), queenless hub workers (dark yellow), and queenless non-hub workers (yellow). Solid lines and shaded bands show the population-level mean and 95% confidence interval predicted by the `glmmTMB` interaction model (Degree *∼* QR_Queen_Condition *×* ovary_idx + (1|Trial)) reported in the main text; individual points show colony-level mean values per bee.

We observed a larger betweenness centrality observed in queens compared to hub bees (Figure S15), likely due to betweenness being a measurement of how well an individual connects disparate groups in a network (Farine and Whitehead, 2015). Because queens interact significantly more than workers, they will necessarily be on the most centralized path between disparate individuals. In queenless colonies, which have multiple queen-like hub bees, no individual bee is guaranteed to be on the shortest path.

Because hub bees display network behavior profiles similar to their natal colony’s queen, we next asked if they also assumed a queen-like reproductive physiology. Indeed, hub bees had a larger ovary index than the non-hub queenless workers, suggesting that these individuals activate their ovaries most strongly in queenless colonies (Figure 4b: *χ*^2^ = 25.4, *p* = 2.88 *\**10^*−*133^). This coincides with a significant interaction between condition and *×* ovary index on interactivity (Degree*∼* QR ovary_idx + (1 | Trial), glmmTMB: *n* = 264, interaction *β* = 10541, *z* = 6.42, *p* = 1.37 *×*10^*−*10^), indicating that the slope of interactivity on ovary index differs between queenright and queenless colonies. Specifically, queenless workers showed a significant positive relationship between interactivity and ovary index (emtrends slope = 8843, SE = 1072), while queenright workers did not (slope =*−* 1698, SE = 1281; pairwise slope comparison: *t* = *−*6.42, *p <* 0.0001) (Figure 4c). This surprised us, as no physiological measures were considered in our behavioral PCA at all. Importantly, this lack of behavioral variation in queenright settings is not the result of a lack of physiological variation – there is already variation in ovary size between workers in these queenless colonies (Figure 4c). As a result, this suggests worker social network position is highly predictive of reproductive status and *vice versa* in queenless, but not queenright, contexts.

### Latent Behavioral Variation Among Workers Reshapes Bumble Bee Social Network Structure

We next asked if the increase in interactivity of queenless partitions was driven by interactions between hub bees, analogous to dueling tournaments in queenless ponerine ants (Opachaloemphan et al., 2021). To examine this, we calculated assortativity between hub bees by testing how enriched hub-to-hub interactions were relative to hub-to-non-hub interactions. We found no evidence for increased assortativity between hub bees (Figure S16), suggesting consistent mixing of queenless workers regardless of their behavioral profile.

Taken together, our results suggest that hub bees in queenless environments express a specific behavioral syndrome in which they are more interactive and more locomotive, yet also more spatially restricted. It is important to note that we cannot disentangle causality between these behavioral phenotypes: bees moving faster will contact one another more frequently, sometimes antennating in the process, but bees may also be moving faster because it allows for an increase in antennation rate.

Finally, we asked whether the presence or absence of the queen was associated with differences in colony-level network structure, particularly in relation to the emergence of behaviorally distinct workers. While queenright partitions were highly centralized around a single queen, queenless colonies contained multiple, highly connected hub bees (Figure 5a). These structural differences coincided with the expression of latent behavioral variation among workers and a change in overall network properties. This unmasked variation was consequential for how information transfer occurred in each partition. Transitivity, a measure of “cliquishness” (subgroups formed within a social network) (Table S1), was higher in queenless networks for four of five replicates (Figure 5b: *χ*^2^ = 380, *p* = 1.19 *\**10^*−*84^). We next explored global efficiency, a measure of information flow that tracks the shortest path length between all pairs of nodes within a network (Latora and Marchiori, 2001). Queenless networks were more efficient, (Figure 5c: *χ*^2^ = 1311, *p* = 3.64*\** 10^*−*287^), suggesting that the larger total number of interactions in queenless colonies (Figure 2a: *χ*^2^ = 1619, *p ≈*0) may facilitate a more rapid spread of information. Finally, queenless colonies were less disassortative than their queenright counterparts (Figure 5d: *χ*^2^ = 121, *p* = 3.74 *\**10^*−*28^), meaning the tendency of individuals with dissimilar numbers of connections (i.e., degree) to interact was weaker among queenless nestmates. This reflects a shift towards a more neutral pattern of interactions among workers (i.e., neither an assortative nor disassortative network). Together, these findings suggest that the emergence of hub bees in the queenless context is associated with a broader restructuring of level-colony social networks, with network properties consistent with more distributed connectivity and potentially faster spread of interactions..

**Figure 5.**
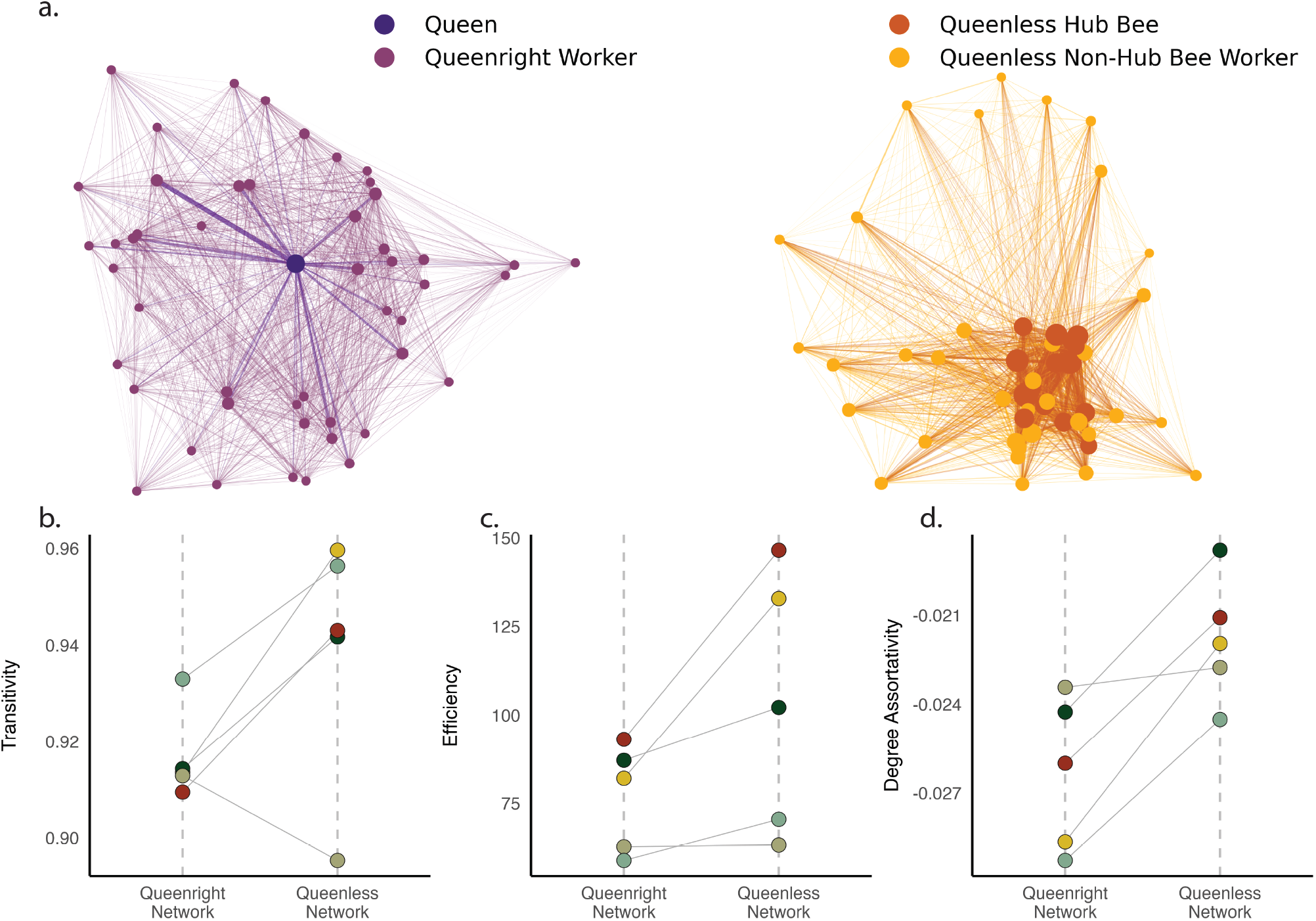
Network properties in queenless colonies are consistent with enhanced efficiency. Network plots of queenright and queenless colony example (Colony 4) and network metrics plotted by colony status. (a) (left) Colony 4 queenright network weighted by pairwise interaction count. (right) Colony 4 queenless network weighted by pairwise interaction count. (b) Network transitivity by queenright/queenless status. (c) Network efficiency by queenright/queenless status. (d) Degree assortativity by queenright/queenless status.

## Discussion

Social insect colonies are often organized around a dominant reproductive queen whose presence impacts reproductive hierarchies and colony social organization. Yet how social structure can be reorganized or maintained in her absence is less well understood. Here, we used matched queenright and queenless partitions of bumble bee colonies to examine how changes in social context impact both individual behavior and group-level organization. By tracking over 80 million pair-wise interactions among individually tagged workers, we found that behavioral variation was higher in queenless colonies compared to their matched queenright parti-tions. A strong association between behavioral variation and reproductive physiology emerged in queenless conditions that was not observed in the presence of a queen. In queenright colonies, the queen herself occupies this central network position. Together, this indicates that reproductive physiology predicts network centrality across both social contexts — but the emergence of hub bees among workers is specific to queenless conditions. A subset of queenless workers exhibited behavioral profiles that were more extreme than matched, queenright workers, and the presence of these individuals was associated with network reorganization at the colony level. Together, these findings suggest that latent variation among workers can scale to shape emergent properties of social organization and provide insights into how collective systems can remain flexible and resilient, even in the absence of dominant individuals.

### The Queen Occupies a Central, Yet Passive, Role in the Colony Social Network

Bumble bee colonies are highly interactive, leveraging both physical and chemical communication to coordinate behavior and regulate reproduction (Jandt and Dornhaus, 2011; Padilla et al., 2016; Orlova, Treanore, and Amsalem, 2020). Previous studies have indicated that physical contact between queens and workers is essential for maintaining reproductive division of labor (Padilla et al., 2016). Our findings reveal that queens serve as the most highly connected nodes in queenright interaction networks, but these interactions are driven primarily by worker-initiated contact. Queens occupied a relatively restricted spatial range within the colony, consistent with previous observations of queen localization near brood (Cnaani, Schmid-Hempel, and Schmidt, 2002; Jandt, Huang, and Dornhaus, 2009), suggesting that queen centrality may emerge from worker-driven strategies. The queen’s passive centrality may also reflect her large body size or potential constraints that arise from egg-laying and thermoregulation of developing workers (Helms and Kaspari, 2015).

Aggression is a commonly used strategy to maintain reproductive dominance within many social insects and vertebrate societies (Holekamp and Strauss, 2016; Tibbetts, Pardo-Sanchez, and Weise, 2022). However, unlike the closely related species, *B. terrestris* (Pandey, Motro, and Bloch, 2020), *B. impatiens* queens appear to maintain reproductive dominance through non-aggressive mechanisms. *B. impatiens* queens rarely display overtly aggressive behaviors such as lunging, biting, or stinging (Padilla et al., 2016). This pattern differs markedly from other simple eusocial species that often rely on aggressive enforcement of dominance (Brothers and Michener, 1974; Jandt, Tibbetts, and Toth, 2014) as well as complex eusocial species like honey bees (*Apis mellifera*) where queens maintain their reproductive monopoly through pheromonal con-trol (Winston, 1987). *B. impatiens* queens are substantially larger and express a distinct cuticular chemistry that reflects reproductive status (Orlova and Amsalem, 2021), potentially providing information on dominance status without a need for aggressive reinforcement (Tibbetts, Pardo-Sanchez, and Weise, 2022). For example, workers could be using standard interactions as a way to monitor their reproductive dominance status (Alaux, Jaisson, and Hefetz, 2006) or to advertise their sterility, potentially reducing within-colony conflict (Amsalem, Twele, et al., 2009).

### Latent Variation in Worker Behavior and Physiology Manifests in a Queenless Environment

We observed increased behavioral variation among queenless workers and a small subset of individuals that expressed queen-like behavioral and physiological profiles that were not present in queenright conditions. These “hub bees” exhibited high interactivity and movement while remaining spatially constrained. Hub bees were identified solely based on behavioral metrics in our principal component analysis, but these individuals also displayed increased ovarian development relative to other colony members.

It remains to be determined whether the extreme behaviors of these individuals influence physiological changes or vice versa. Similar to other social insect societies (Sharma, Gadagkar, and Pinter-Wollman, 2022), the reproductive dominance observed in hub bees likely emerges from a constellation of interacting factors. Hub bees tend to be slightly larger than nestmates, but size alone is insufficient to explain their reproductive dominance. Alternatively, hub bees may simply have naturally lower thresholds for the expression of queen-like behaviors when the queen is lost (Beshers and Fewell, 2001).

The coupling between behavior and reproductive physiology in the hub bees appears to be context-specific. Highly interactive workers in queenright partitions did not express similar rates of ovary activation, suggesting that queen presence suppresses the expression of latent behavioral and physiological differences among workers. Future work incorporating temporal comparisons of queen presence and loss can help to determine whether hub bees exist in queenright conditions but remain behaviorally or physiologically suppressed.

### A Subset of Workers Reshapes Queenless Colony Dynamics

Because queenless colonies exhibited greater behavioral variance and a small subset of high-centrality workers, we next asked whether these individual-level changes coincided with predictable shifts in colony-level structure. While queenright colonies were highly centralized around a single queen, queenless colonies developed a decentralized social architecture with multiple, highly connected individuals. This shift coincides with an increased overall interactivity and higher network efficiency, transitivity, and degree assortativity. Taken together, these network metrics suggest that, within our experimental setup, queenless colonies possess enhanced information-sharing capacities.

While these graph metrics are proxies rather than direct measurements of information flow, their joint shift in queenless partitions is consistent with more distributed connectivity and higher potential throughput of interactions. Such shifts in network-level dynamics can carry dramatic implications for the physiology of both the individual and the collective in social insects (Smith, 2018; Bonabeau, Theraulaz, and Deneubourg, 1999; Kay et al., 2024). The transition to a distributed rather than centralized network structure could help to enhance a colony’s ability to fine-tune task allocation (Pradhan, Patra, and Chowdhury, 2021; Fisher et al., 2022), as workers can receive more frequent and diverse social signals through the restructured network. Distributed control and frequencies of physical contact are known to play an important role in task allocation in several ant species, including foraging behaviors (Gordon, 2016; Gordon, 1996). However, increased interactivity and connectivity may also carry a cost: higher interaction rates and the presence of multiple reproductive individuals may impose additional energetic demands on the colony, though the magnitude of such costs remains to be directly measured (Waters, 2014).

## Conclusions

Our work highlights how social context can modulate the expression of individual traits such that collective dynamics are altered. Here, we compared queenright and queenless partitions of bumble bee colonies to link individual-level behavioral traits to reproductive physiology and colony-level shifts in social organization. Our observations show that, in experimental queenless environments, latent behavioral and physiological variation among workers is unmasked, and that the emergence of these new social phenotypes is associated with a shift in colony-level network organization. In *B. impatiens*, queen loss is a natural part of the annual colony cycle, as colonies transition from a cooperative phase with a single, reproductive queen to a competition phase where multiple workers lay male-destined eggs (Goulson, 2010). Notably, despite this transition, overall network structure remained robust: queenless colonies reorganized into distributed, highly connected architectures rather than fragmenting. Latent behavioral variation among workers could serve as a reservoir of variation that enables rapid transitions between organizational states as conditions change (Blight et al., 2016). A key open question is how this latent variation is regulated in the presence of the queen and what mechanisms trigger its expression upon queen loss. Future work combining detailed behavioral tracking with molecular and neurobiological approaches will be essential to determine how queen-derived signals suppress behavioral and physiological differentiation, and how individual-level variation shapes collective outcomes across the colony cycle.

## Methods

### Animal Rearing and Tagging

Our dataset consists of ten 96-hour (4-day) videos from five *Bombus impatiens* source colonies, each split into a queenright and a queenless partition (10 arenas total), recorded continuously under controlled conditions. Our source colonies were obtained from Koppert Biological Systems (Howell, MI USA); no males nor gynes were observed in any source colony, suggesting each colony arrived in its respective cooperative phase. Each colony was maintained in a warm (28^*°*^C), quiet room illuminated by red light (which bees cannot see) to minimize disturbance. Tracking was performed within 10 days of colony arrival to minimize the possibility of stress due to overcrowding, as workers eclose on a daily basis. All experiments were performed between November 11, 2022, and February 7, 2023.

ArUco tags generated from the 5X5_50 set were printed on TerraSlate 5 Mil paper (TerraSlate, Englewood, CO USA), which is waterproof and does not tear, and cut to 4.25*×*4.25 mm using a Silhouette Cameo cutting machine (Silhouette, Lindon, UT USA). On the morning of each tracking experiment, a single queenright colony was moved to a 4^*°*^C room to reduce handling stress when the colony was opened and bees removed. Size-matched groups of bees were gently removed with soft forceps and placed in conical vials submerged in wet ice only until no movement was observed. Next, individual bees were removed and a small drop of cyanoacrylate glue (Loctite, Hartford, CT USA) was placed on the dorsal thorax, to which a single ArUco tag was applied using the tip of a pin. Our tagging strategy caused minimal apparent stress as bees actively rewarmed and resumed normal activities within minutes after arena placement. Cyanoacrylate glue does not affect behavior or mortality in bees (Gernat et al., 2018) and previous work only observed an acute increase in grooming following a similar tagging strategy (Crall, Gravish, Mountcastle, and Combes, 2015).

Tagged bees were placed in one of two 27 x 27 cm laser-cut arenas that were designated as “queen-less” or “queenright,” and the colony’s queen was tagged and added to the latter arena (Figure 1). Each partition contained 48-52 workers. Arenas contained silicon matting, on which bees can easily walk, and 5g of brood from the source colony. Four cotton wicks soaked in nectar substitute (equal parts pure sugar water and inverted sugar water with added feeding stimulant and amino acid supplementation) were placed in one corner of the arena to allow *ad libitum* feeding, and 5g of ground honey bee pollen mixed with nectar substitute at a ratio of 10:1 (pollen:nectar substitute) was added for protein nutrition and a more naturalistic environment. Temperature was maintained between 27 and 29^*°*^C.

We note that our experimental setup used two-dimensional arenas without a separate foraging chamber, which may affect the range of behaviors expressed. However, bumble bees, unlike honey bees, do not need to leave the hive to perform foraging or cleansing flights, a feature leveraged in this study as out-of-hive conditions like resource availability, weather patterns, or physical trauma from flight could not affect our results. Moreover, similar enclosed-arena designs have been widely used to study bumble bee social behavior and have demonstrated that key colony-level behaviors persist under these conditions (Crall, Gravish, Mountcastle, Kocher, et al., 2018; Pandey, Motro, and Bloch, 2020; Wolf et al., 2023).

### Tracking

Arenas were covered with clear acrylic and lit using 7 high-intensity 850nm LED light bars (Smart Vision Lights L300 Linear Light Bar, Norton Shores, MI USA) to allow continuous imaging without disturbance. 5 hours after establishing the partitioned colonies, we imaged the arenas from above using a Basler acA5472-17um (Basler AG, Ahrensburg, Germany) camera recording 3664px *×* 3664px frames at 20 frames per second (14.5 for Colony 2). Recordings were taken using a modified version of CAMPY, a Python package developed for real-time video compression (Severson, 2021). The resulting videos have a spatial resolution of *∼*15.5 pixels/mm, allowing us to capture fine-grained behaviors. Each bumble bee worker varies between approximately 9mm and 14mm in length, so the resulting pixel length of each worker is approximately 139.5px to 217px in the video data (Williams et al., 2014).

After video acquisition, we utilized SLEAP to capture bee pose. We used a 9-node skeleton marking the head (mandible), two thorax points, abdomen, left and right antennal joints, left and right antennae tips, left and right wings, the pretarsus of each leg, and the ArUco tag. We trained two separate models, one for the workers and one for queens to appropriately account for morphological differences. For workers, we trained on 107 frames resulting in a final mean error distance across all nodes of 7.11px (0.46mm) in our validation set. For the queen model, we trained 361 frames and the resulting model has a mean error distance across all nodes of 30.09px (1.94mm). All models are provided in Data Availability.

We utilized NAPS, which integrates SLEAP and ArUco, to ascertain individual bee identities after estimating pose. Using the tracklets identified by SLEAP, a Hungarian matching algorithm was employed to resolve identification ambiguities using the ArUco tags, as described in (Wolf et al., 2023). Post-identification, we filtered data for five anomalies: ‘Tag Identity’ ‘Jumps’, ‘Between Jumps’, ‘Skeleton Irregularities’, and ‘Spatial Irregularities’:

1. We removed all nodes that were mapped with SLEAP to tags not used in the experiment.Jumps were defined as instances where between two frames a node either (i) moved greater than 10 pixels (*∼*0.65 mm), when fewer than 80 percent of other nodes moved less than other nodes (node jump) or (ii) when greater than 80 percent of other nodes moved greater than 100 pixels (*∼* 6.5 mm) (tag jump).
2. Between jumps were regions within 10 frames of a tag jump on both directions. In this case we removed all nodes.
3. Skeleton irregularities were regions where length of the edge of a skeleton was greater than 5 z-scores above the mean. In this case we removed both connection nodes.
4. Spatial irregularities were regions where two nodes on the same bee were less than 2 pixels away from each other (typically a result of the same node being indicated twice).

The mean filtering of each step of this data is in Supplementary Figure 3.

### Touch Detection

We aimed to quantify the modalities of bumble bee contacts, particularly focusing on head-to-head antennation as an indicator of social interaction. To do this, we converted the skeleton into regions of the body by mapping the space a buffer distance B from the skeleton Figure 1 in cyan). This allows us to define physical interactions as any frame with overlapping regions between two bees. Head-head interactions are defined by overlaps between two antennal regions, and Head-body interactions are defined by overlaps between an antennal region and a thorax/abdomen region. To appropriately standardize our data, we calculated the area and perimeter of our regions. The skeleton used to map interactions can be found in Figure S1 and Data Availability.

### Network Analysis

Interaction instances were translated into an undirected weighted network. Network data was generated for each hour of our 96-hour recording. Network analyses were conducted using *networkx* version 3.1. We defined weights in two ways: “total Interactions” is the number of unique instances of “bouts” of interaction (lasting at least 2 frames and at least 20 frames away from any other identified interaction between the same two bees). “Interaction Time” is the bouts weighted by their duration such that longer bouts contribute more than shorter bouts.

Using our weighted networks, we calculated both individual-level variables and network level variables as described in (Table S1). These variables were collectively used to create a principal component analysis (Figure S10). We used two different statistical methods to identify the significance of variation in these variables:

### Linear Mixed Models

To identify the impact of worker/queen status on individual- and colony-level network features, we fit linear mixed models with the following model formula:

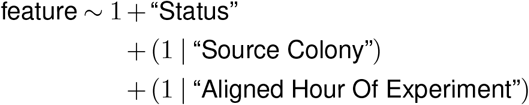

Status (queenright vs. queenless, queen vs. worker, or hub bee vs. non-hub bee, depending on the analysis) is included as a fixed effect, while source colony and experiment hour are included as random effects. Where this study reports both treatment-mean and treatment-variance comparisons on the same response—specifically the colony-level network parameters in Figure 5 and the hub-vs-non-hub comparisons in Figure S14—both p-values are obtained from a single `glmmTMB` (Brooks et al., 2017) model with a Gaussian family and a dispersion sub-model (dispformula*∼* Status) that allows the residual variance to differ between treatments. The Wald *χ*^2^ test on the Status fixed effect provides the mean comparison; pairwise contrasts on the log-dispersion component, computed with the `emmeans` R package (Lenth, 2024), provide the variance comparison. Reporting both effects from the same model ensures that standard errors on the means correctly absorb treatment-specific residual variance. For analyses that report means only (queens vs. workers in Figure 2; queens vs. hub bees in the Discussion), we fit the same fixed-/random-effect structure with the `lme4` R package (Bates et al., 2015) and report Wald *χ*^2^ tests on the fixed effect.

### Velocity

To distinguish between actual movement and apparent movement which occurs as a result of noise in the SLEAP model, we created a histogram of the velocities of each individual. We identified two bimodal peaks representing real movement and subpixel movement, the latter of which we considered to be statistical noise. We used a bimodal distribution (in the method of Crall, Gravish, Mountcastle, Kocher, et al., 2018) to calculate a threshold cutoff for movement for each colony, and calculated both the percentage of time each bee is moving and the average velocity of each bee when moving. Across all workers, mean velocity when moving was *∼*4.8 pixels/frame (*∼*0.31 mm/frame), and workers moved an average of *∼*56% of the time. The 10-pixel node jump threshold corresponds to approximately 2*×* the mean velocity, meaning it prevents the capture of tracking artifacts while retaining the range of normal locomotor behaviors, including rapid approaches. We note that our interaction detection is based on spatial overlap of body regions rather than velocity, so brief, high-speed approach events (e.g., darting) that result in antennal contact would still be captured as interactions even if the preceding movement was flagged as a jump (Amsalem and Hefetz, 2010).

### Directed Network

Insect networks are commonly based on directed dyadic interactions (Appleby, 1983). To identify directionality within our undirected interactions, we isolated the first frame of each bout of interaction and marked the interaction as “directed” if one of the bees in the interaction traveled at least 65.8 pixels (0.5 bee-lengths) more than the other bee in the second before the interaction. From there, our directed network consists of all the directed interactions, with the initiator being the approaching bee and the receiver being the approached bee. Our undirected network includes both directed and undirected interactions, with no differentiation between initiator and receiver.

### Ovary and Body Size Quantification

Ovary dissections were performed at room temperature, and frozen abdomens were allowed to completely thaw in phosphate-buffered saline (PBS) before the tergites T2-T4 were carefully removed. Ovaries were gently lifted out of the abdominal cavity, placed in a new droplet of PBS, and photographed with a Nikon SMZ1270 stereo microscope (Nikon, Tokyo, Japan). The largest oocyte of each ovary was measured in FIJI (Schindelin et al., 2012) and we used the average width across ovaries as a surrogate for reproductive status (Simons and Smith, 2018).

Unlike for honey bees (*Apis mellifera*), bumble bees express size polymorphism that must be accounted for when comparing individuals. Head capsule width, intertegular span, and marginal cell length have all been implemented as proxies for body size (Shpigler et al., 2014; Hagen-Kissling and Dupont, 2013). We measured these structures from *∼*50 workers from four colonies not included in the tracking experiments to show that all three measurements are strongly correlated with body size (Figure S17, Figure S13). While each structure is therefore similarly informative in estimating size variation, marginal cell length offers three major advantages: 1) it is the most easily measured due to lack of curvature or hair present in the head and thorax, respectively, 2) handling the wings does not risk degradation of the body, and 3) mounting wings under clear tape for imaging generates a permanent tissue archive that is easily stored. Finally, dividing average oocyte width by marginal cell size provided an ovary index, a unitless, normalized measure of ovary activation that can be compared across individual bumble (Shpigler et al., 2014; Cini, Meconcelli, and Cervo, 2013). Because measurements could not be easily taken before or during the experiment, we measured marginal cell size for each queenright and queenless worker for a single colony and found there to be no significant difference in size distribution across partitions (Welch’s t-test, t = 1.24, *p* = 0.219), suggesting our visual size-matching was accurately performed.

## Code and Documentation

The analysis code utilized in this study, including scripts for data analysis, visualization, and the generation of network metrics, is publicly available for review and replication purposes. Detailed documentation and source code can be accessed through the following GitHub repositories:

- For the primary analysis code related to this study, please visit: https://github.com/kocherlab/queenright-queenless-analysis.
- For code pertaining to tracking using SLEAP and NAPS, post-processing of tracks, and the computation of network metrics, please refer to: github.com/itraniello/socioQC.

These repositories contain all necessary information for reproducing the tracking, analysis, and network metrics calculations detailed in our study.

## Data Availability

The data utilized in this manuscript will all be available through Princeton’s DataSpace upon acceptance for publication. Paired with code provided in Code and Documentation, this will provide the code and data required to reproduce this work.

## Acknowledgements

This work was supported in part by an NIH Director’s New Innovator Award to SDK (1DP2GM137424-01), the Packard Foundation, the Princeton Catalysis Initiative, and the Department of Physics Undergraduate Research Fund at Princeton University. This work was also supported by the National Science Foundation through the Center for the Physics of Biological Function (PHY-1734030). DMR and SWW were supported by the NSF Graduate Research Fellowship Program (DGE-2039656), IMT is supported by the Lewis-Sigler Scholars Program at Princeton University. The authors thank members of the Kocher Lab for helpful feedback that improved the manuscript.

## Authors’ Contributions

DMR, SWW, IMT, and SDK conceptualized the project; DMR, SWW, AEW, ESW, MLW, DM, and IMT performed formal analyses; DMR and SWW drafted the initial manuscript. IMT and SDK supervised the project. All authors edited the manuscript and provided critical feedback. All authors approved the publication.

## Supplementary Information

**Table S1.**
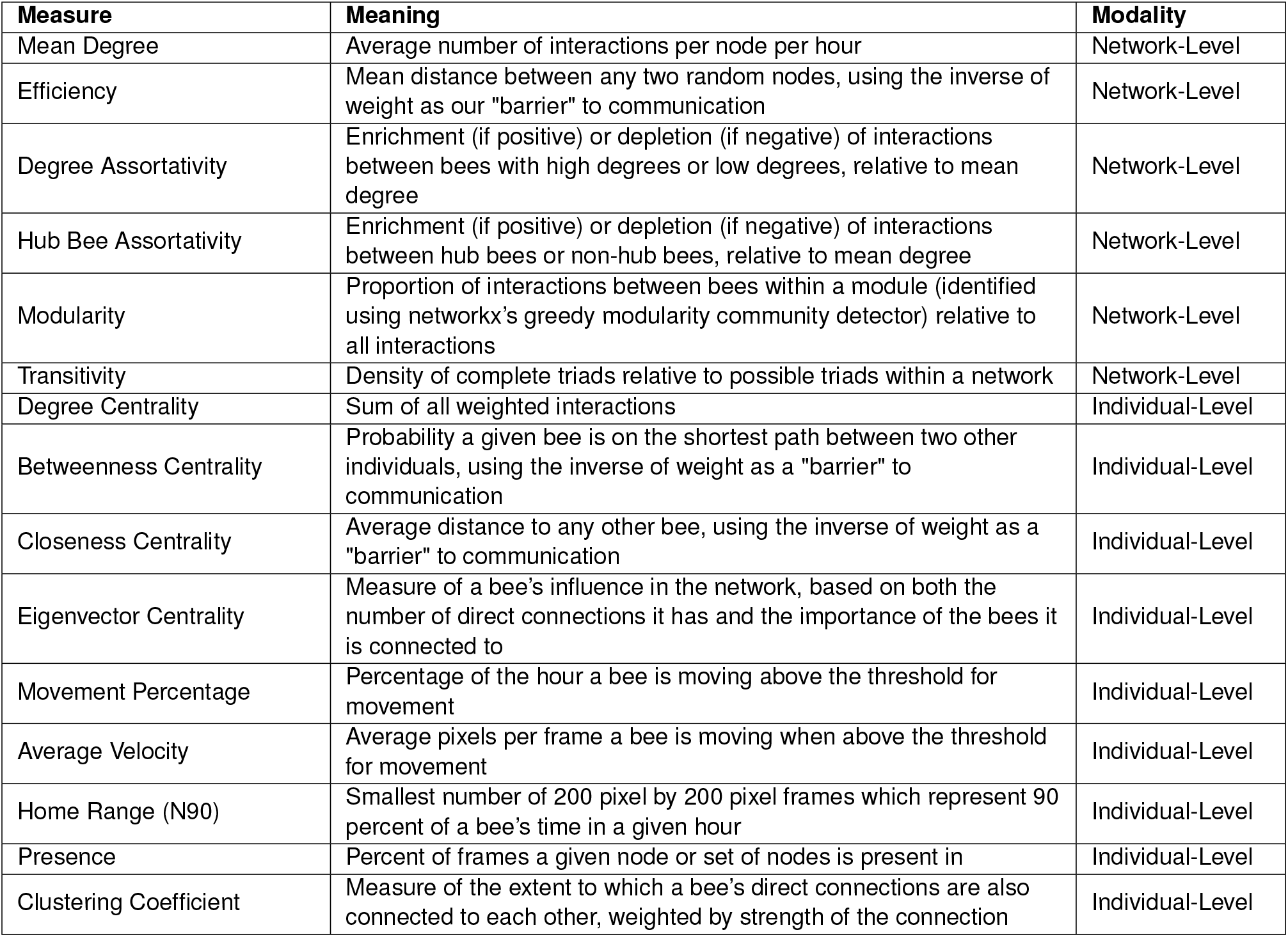
Individual and network-level parameters measured in this experiment.

**Figure S1.**
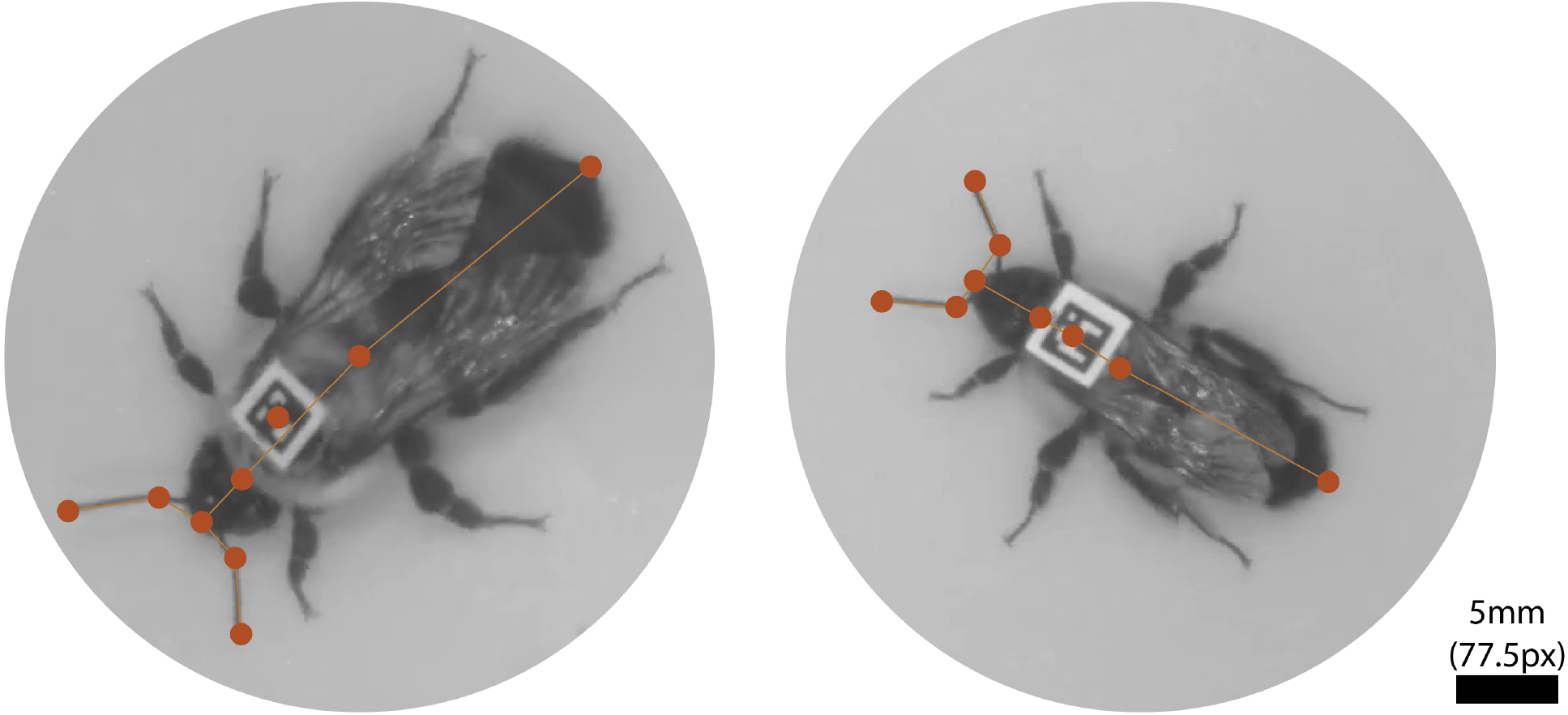
Example SLEAP skeletons on queen and worker showing the nine nodes used for analysis.

**Figure S2.**
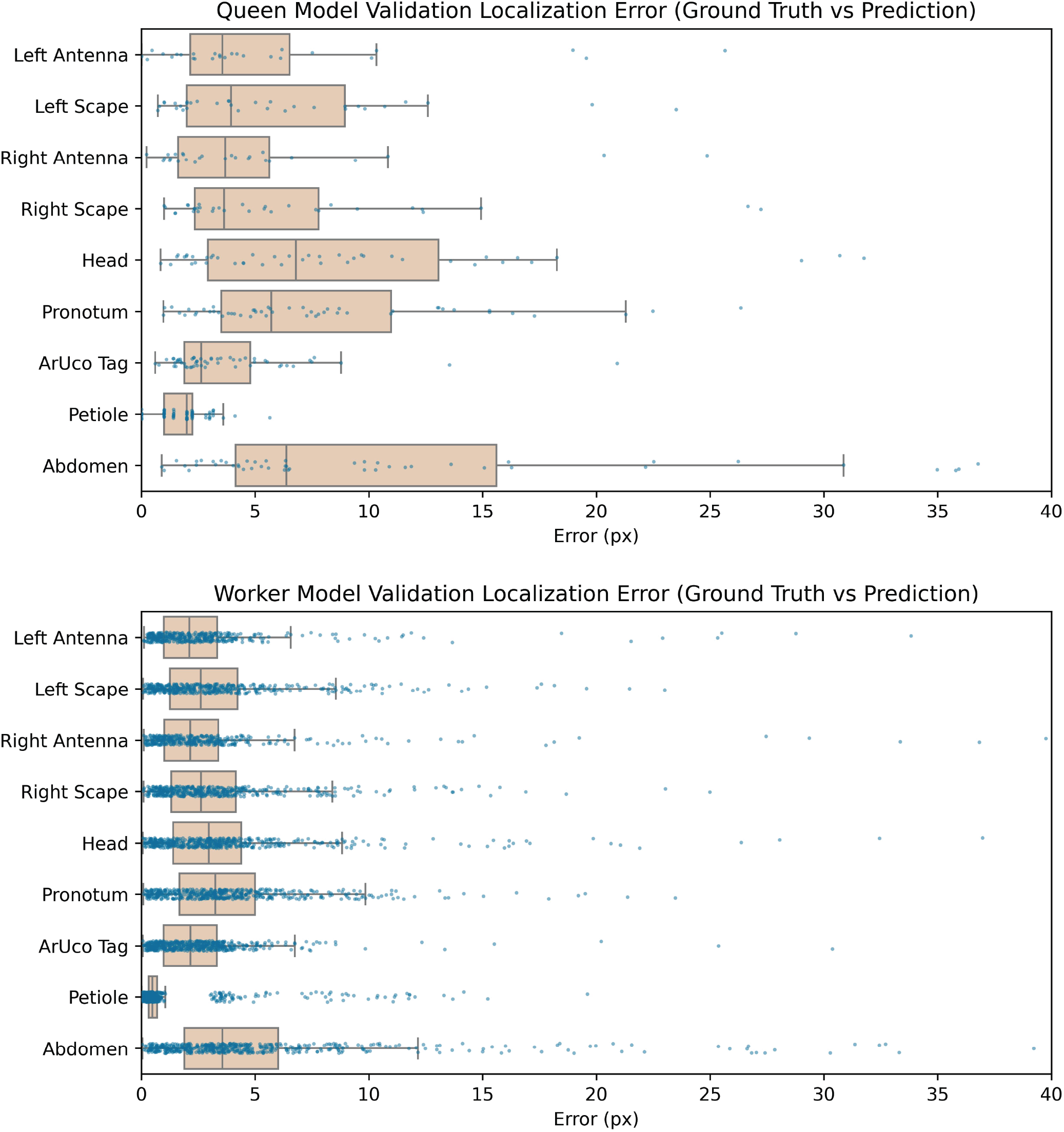
SLEAP model metrics for queen (top) and worker (bottom) models. For this study, px/mm ratio = 15.5, meaning that nearly all localization errors are at the sub-millimeter level. Box and whiskers plots show median, interquartile range (IQR), and upper and lower limits representing 1.5 * IQR above or below the upper and lower quartile, respectively. Each dot represents an instance of detection error in a validation frame.

**Figure S3.**
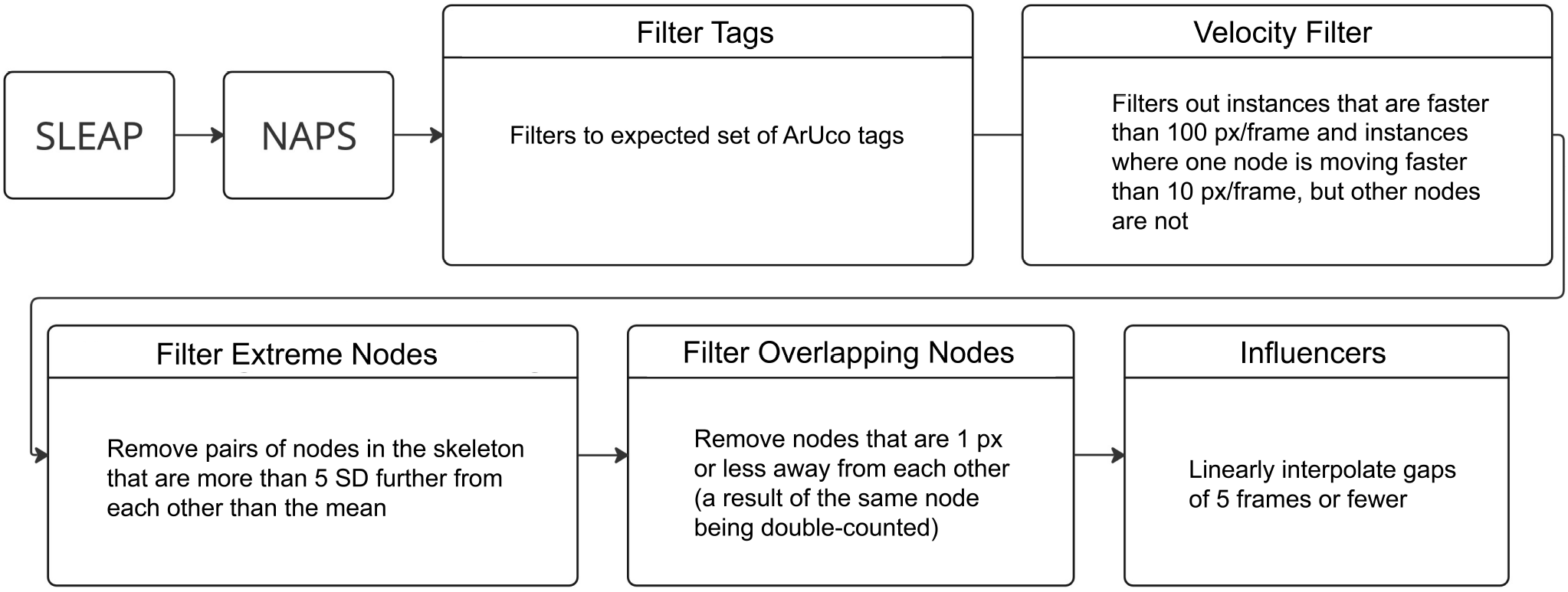
Diagram of filtering steps applied after generating NAPS tracks.

**Figure S4.**
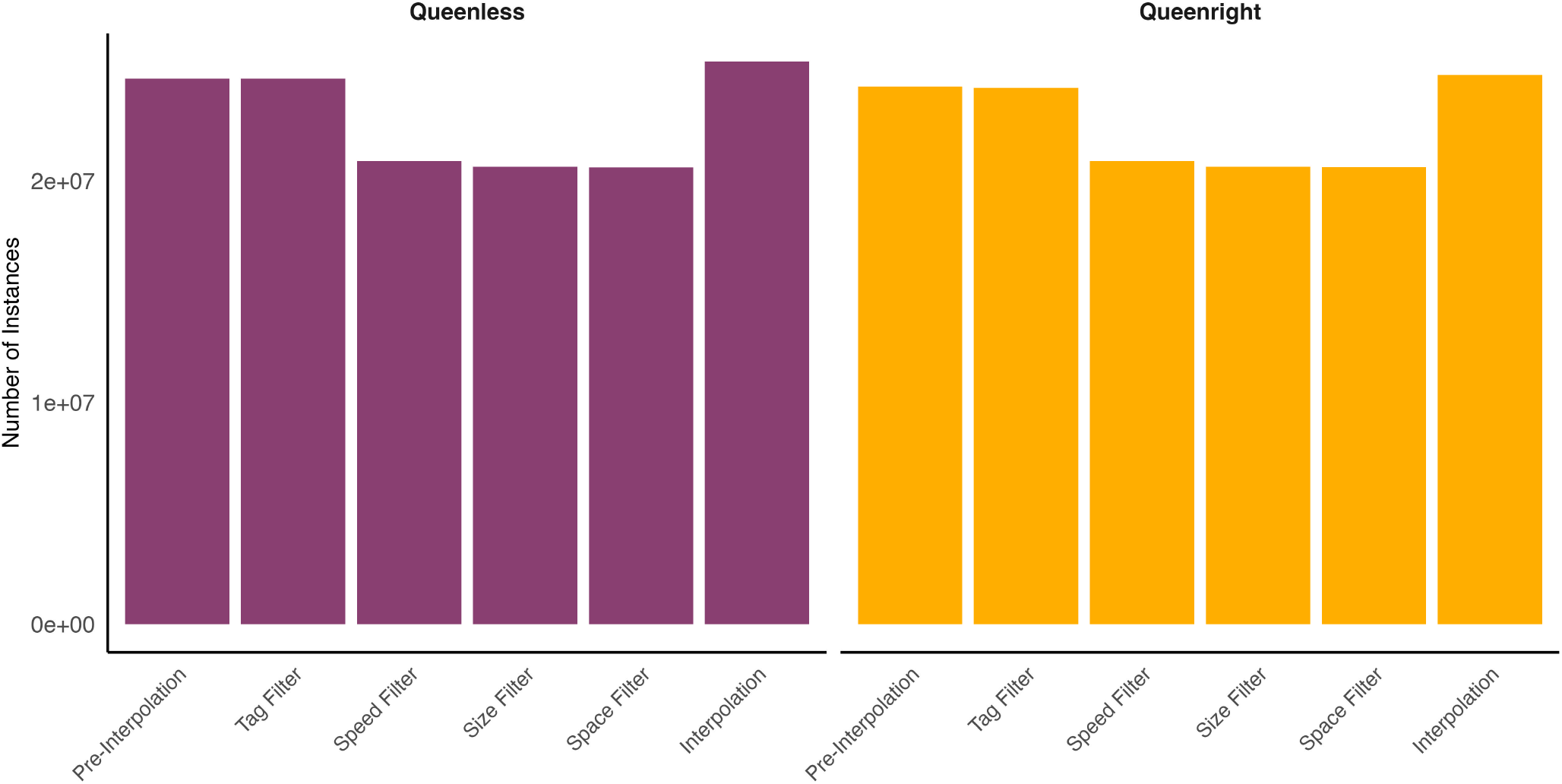
Post-filtering bar plot show total number of instances across all bees across all frames during filtering and interpolation steps.

**Figure S5.**
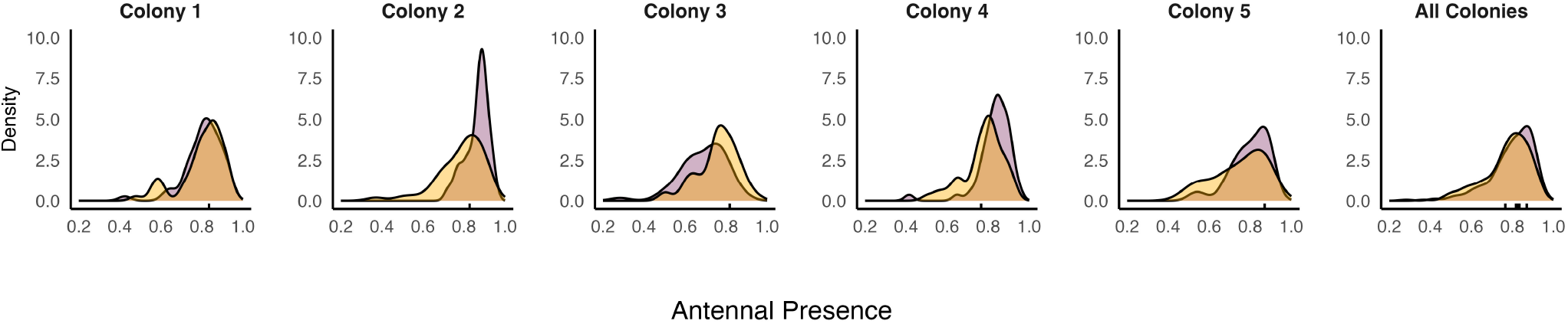
Density plots showing antennal presence per colony.

**Figure S6.**
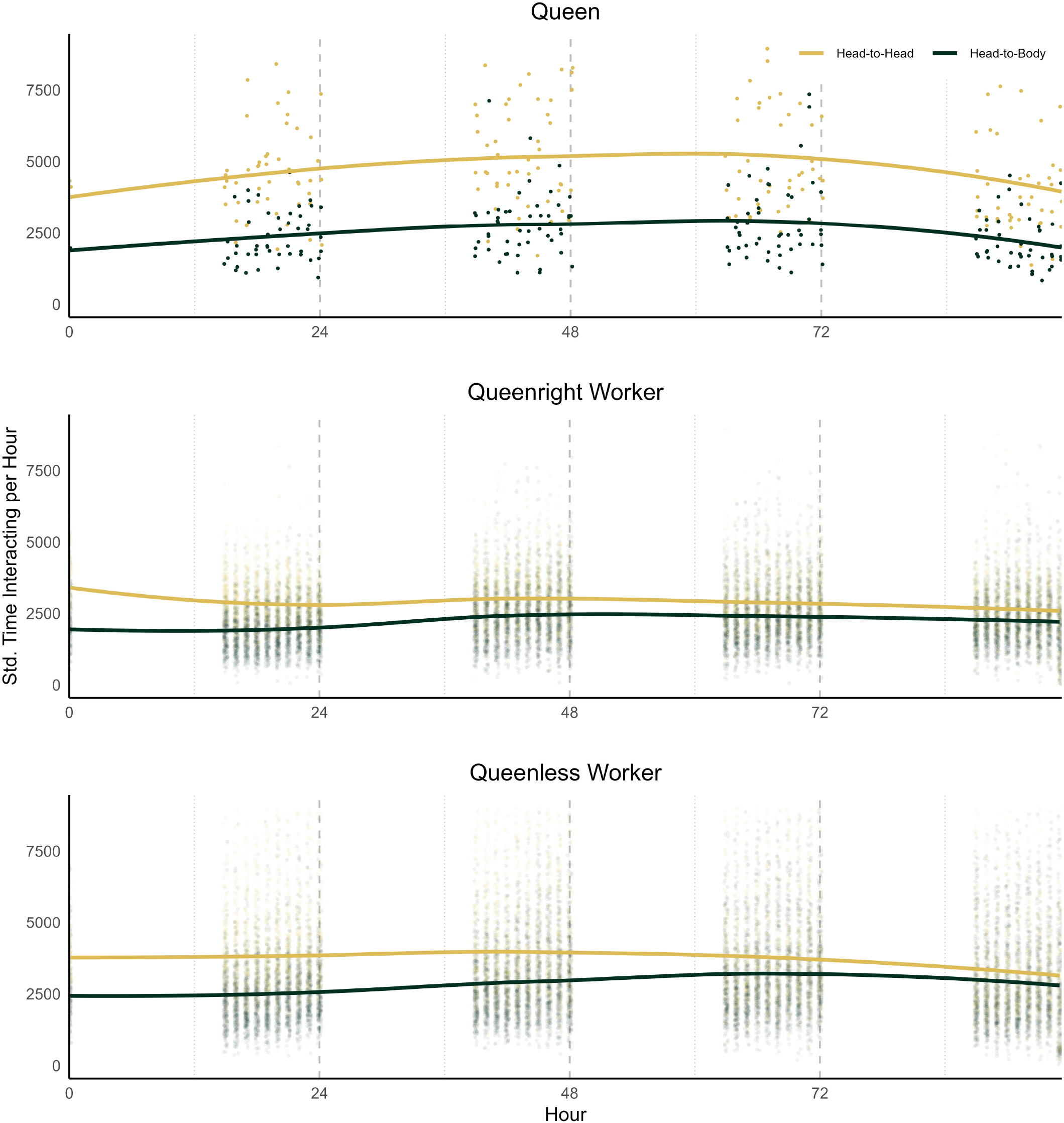
Plots showing standardized interacting time per hour for head-to-head and head-to-body across queens, queenright workers, and queenless workers from 9:00-17:00 each day.

**Figure S7.**
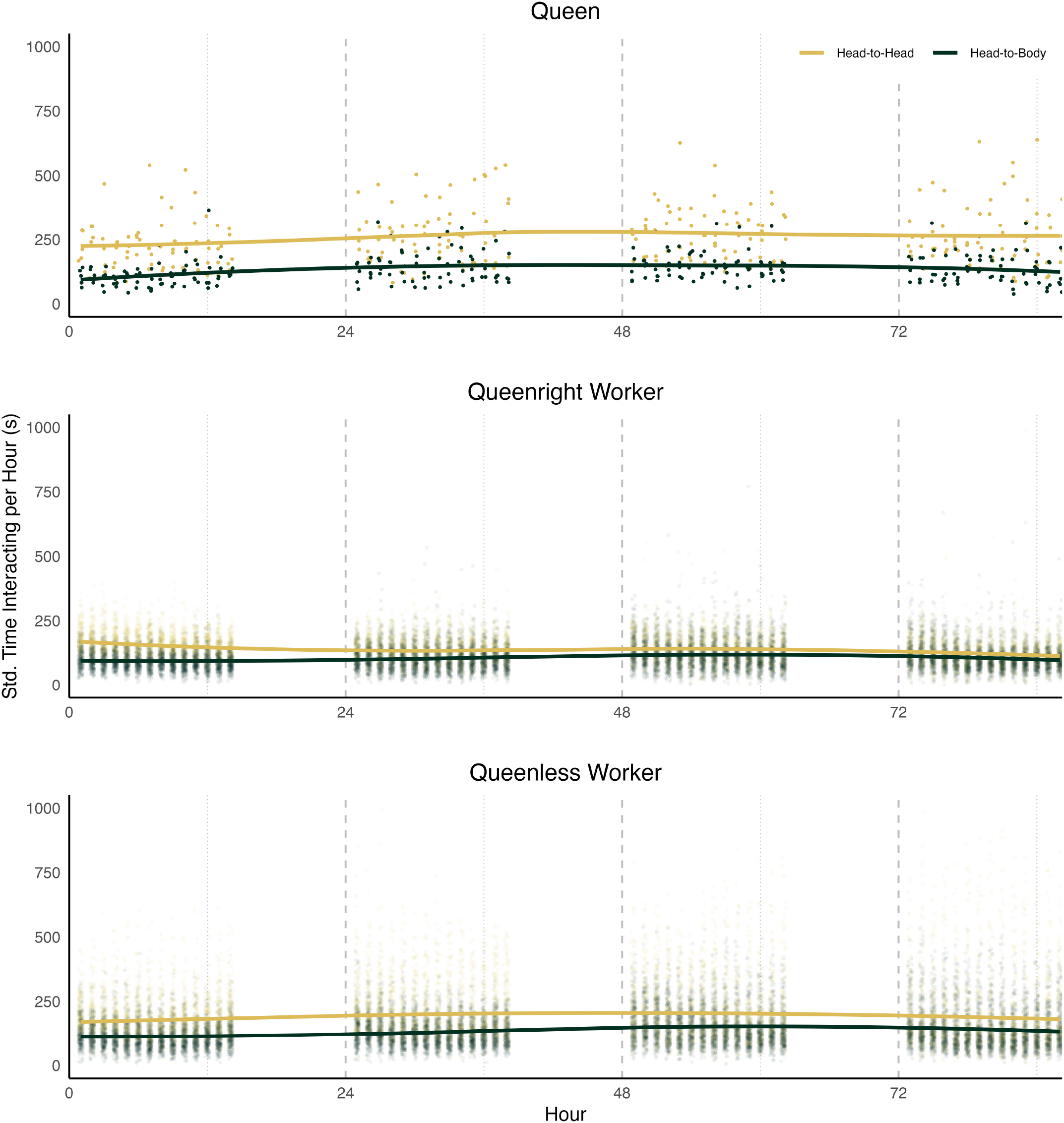
Plots showing status (queen, queenright worker, queenless worker) by interactions per hour during nighttime hours

**Figure S8.**
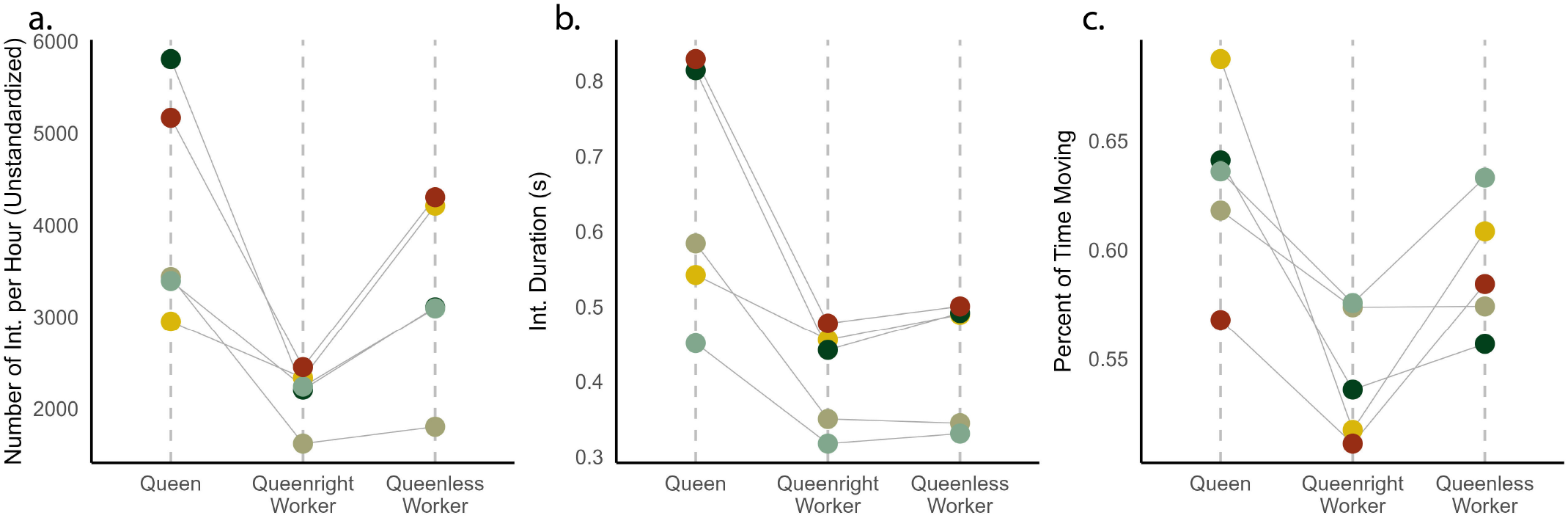
Plots showing status (queen, queenright worker, queenless worker) by individual features: interactions per hour (not standardized to antennal presence), interaction duration, and percent of time moving.

**Figure S9.**
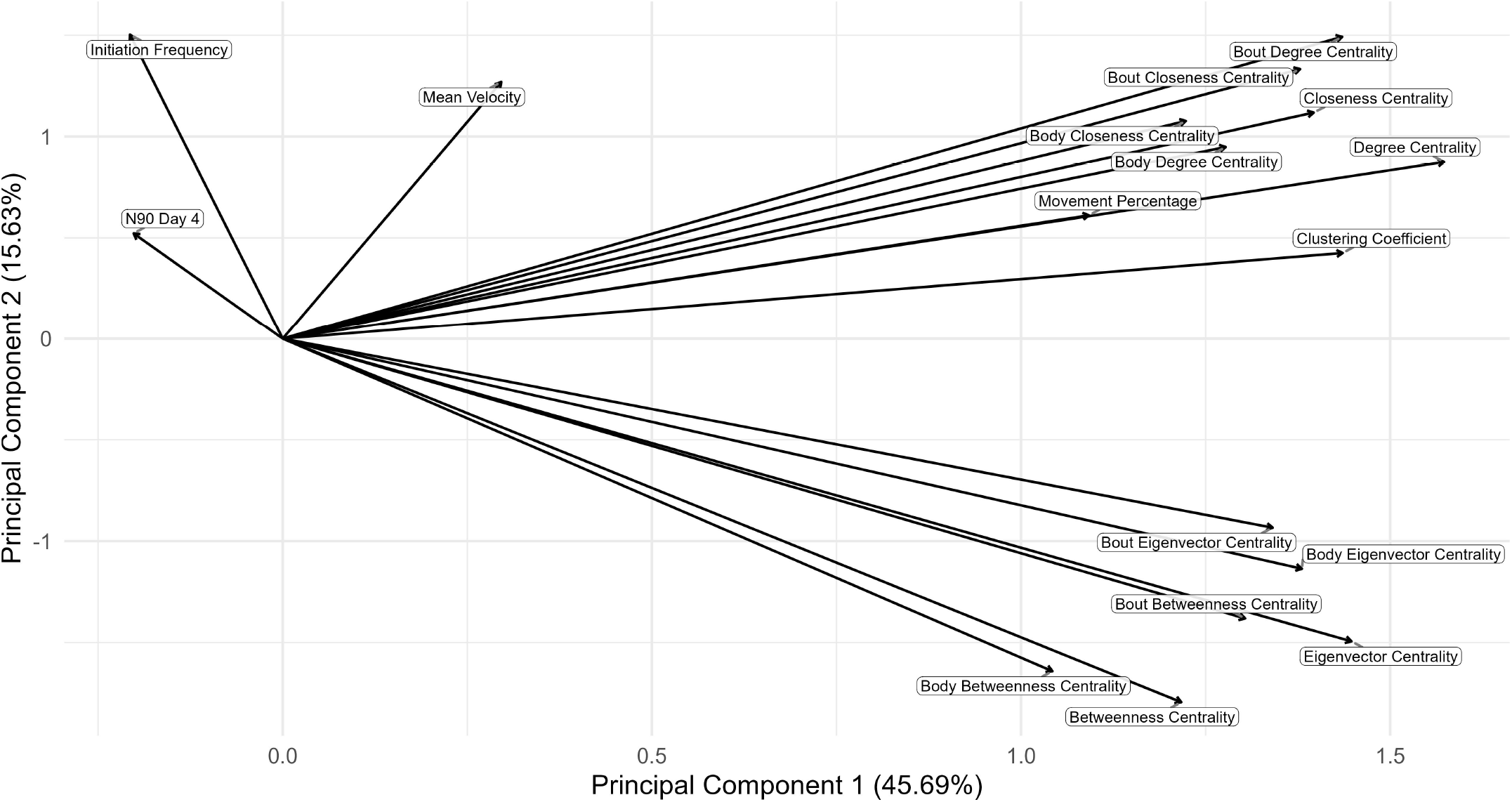
Loading plot showing features used to classify hub bees.

**Figure S10.**
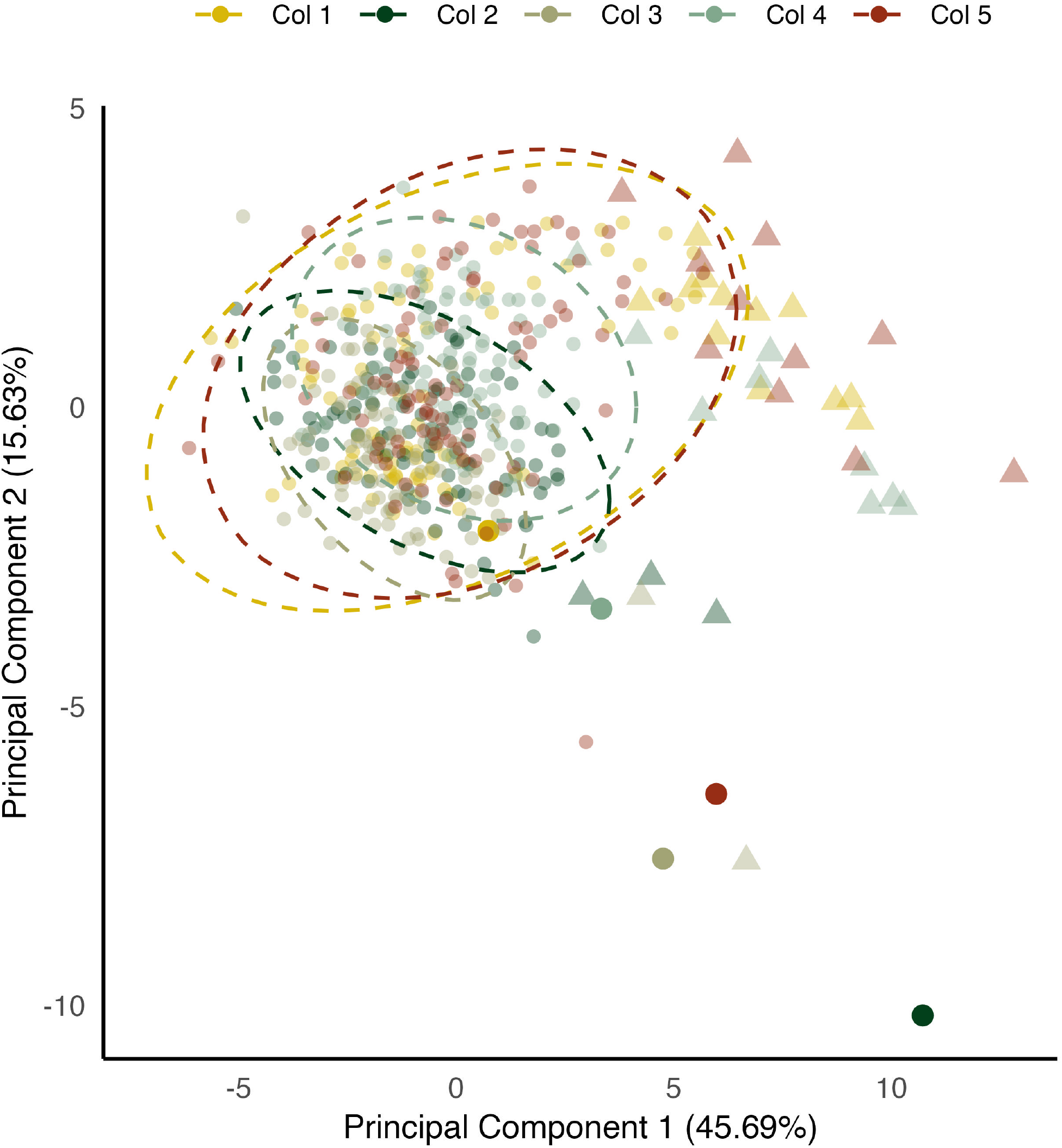
PCA plot from Figure 3a colored by colony with queens enlarged. Outlier bees (triangles) were also enlarged

**Figure S11.**
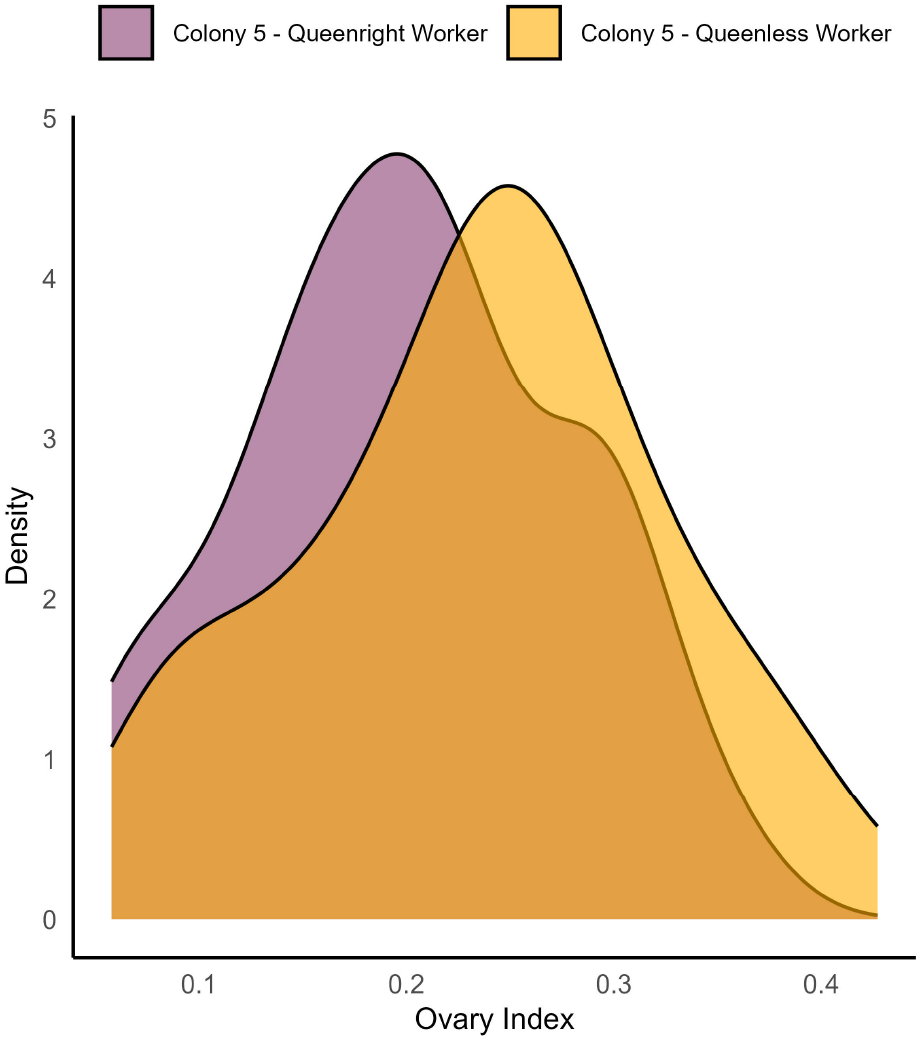
Density plot showing ovary index distributions for Colony 5 grouped by queenright, queenless status. We dissected all individuals in a single colony, Colony 5, to validate our initial hypothesis about variation in ovary index.

**Figure S12.**
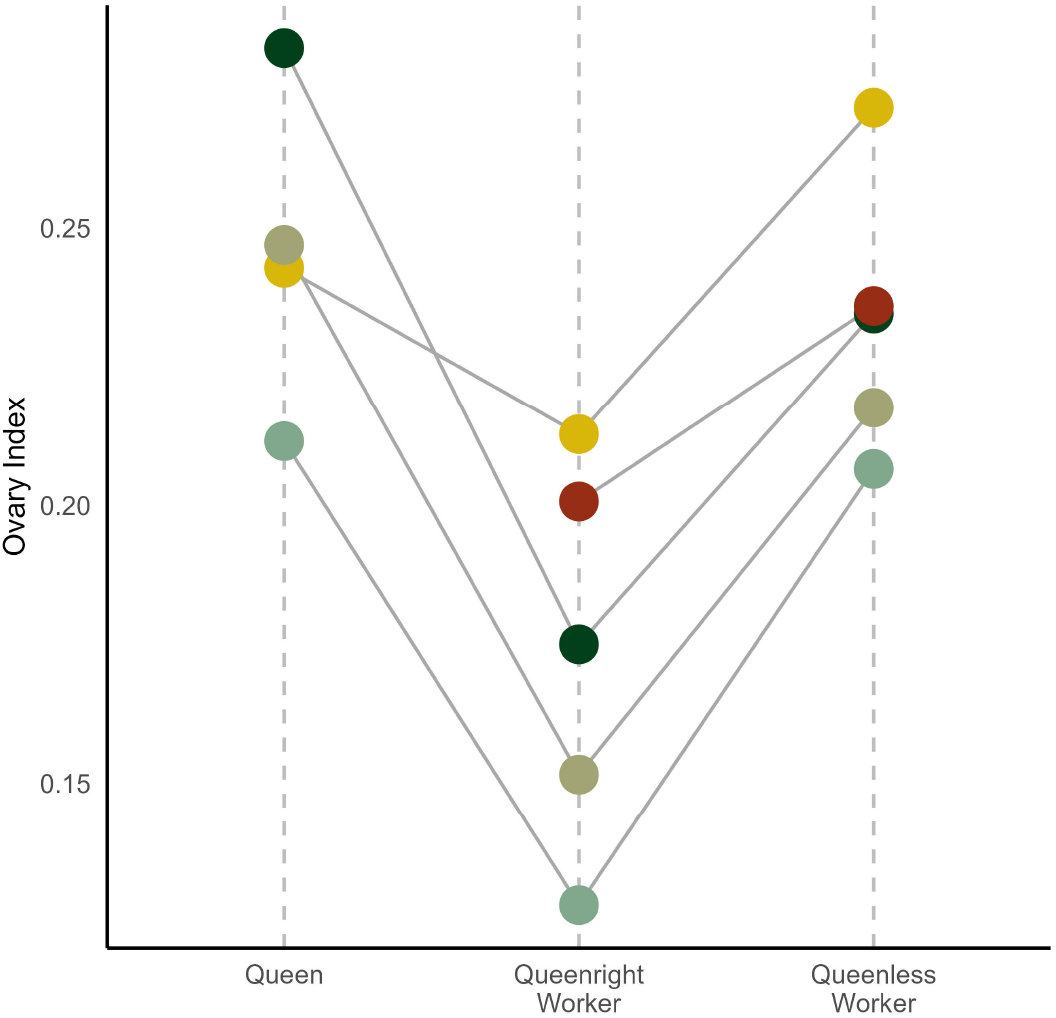
Mean ovary index (width of longest ovariole divided by marginal cell length) by queen, queenright worker, queenless worker status colored by colony.

**Figure S13.**
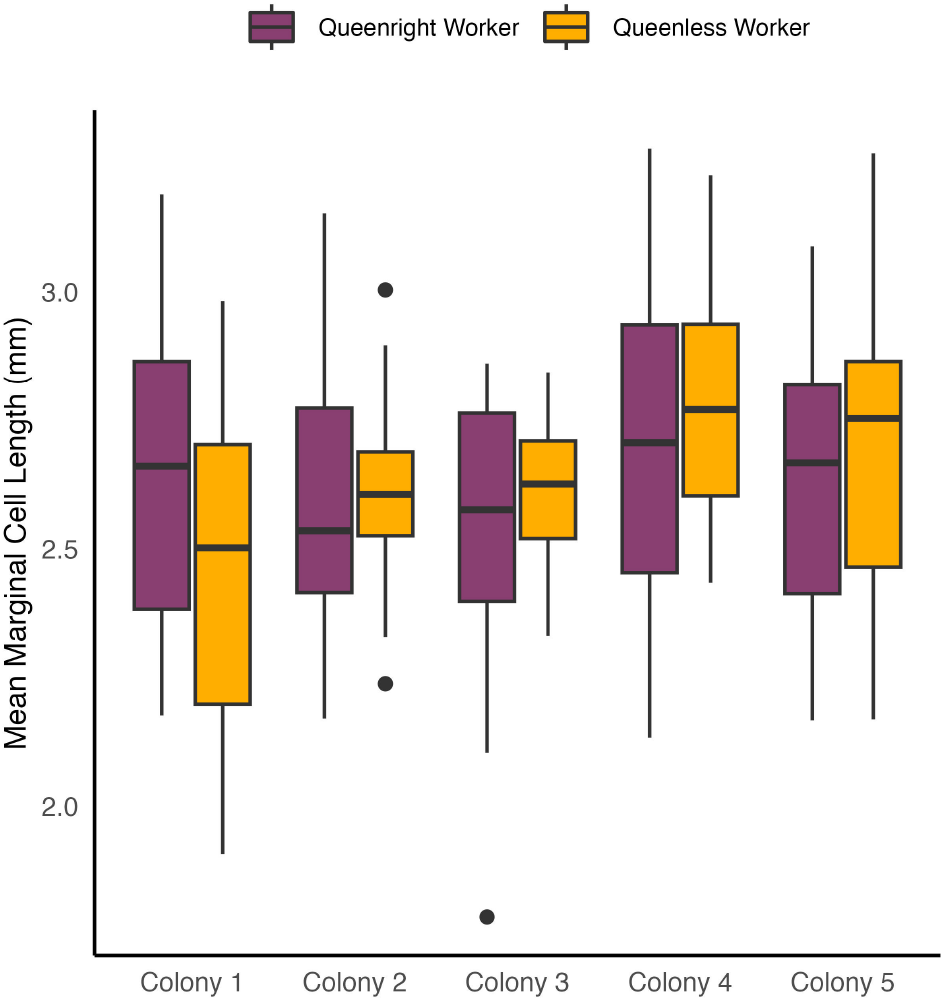
Box plots showing mean marginal cell lengths by queenright (purple)/ queenless (orange) status and source colony.

**Figure S14.**
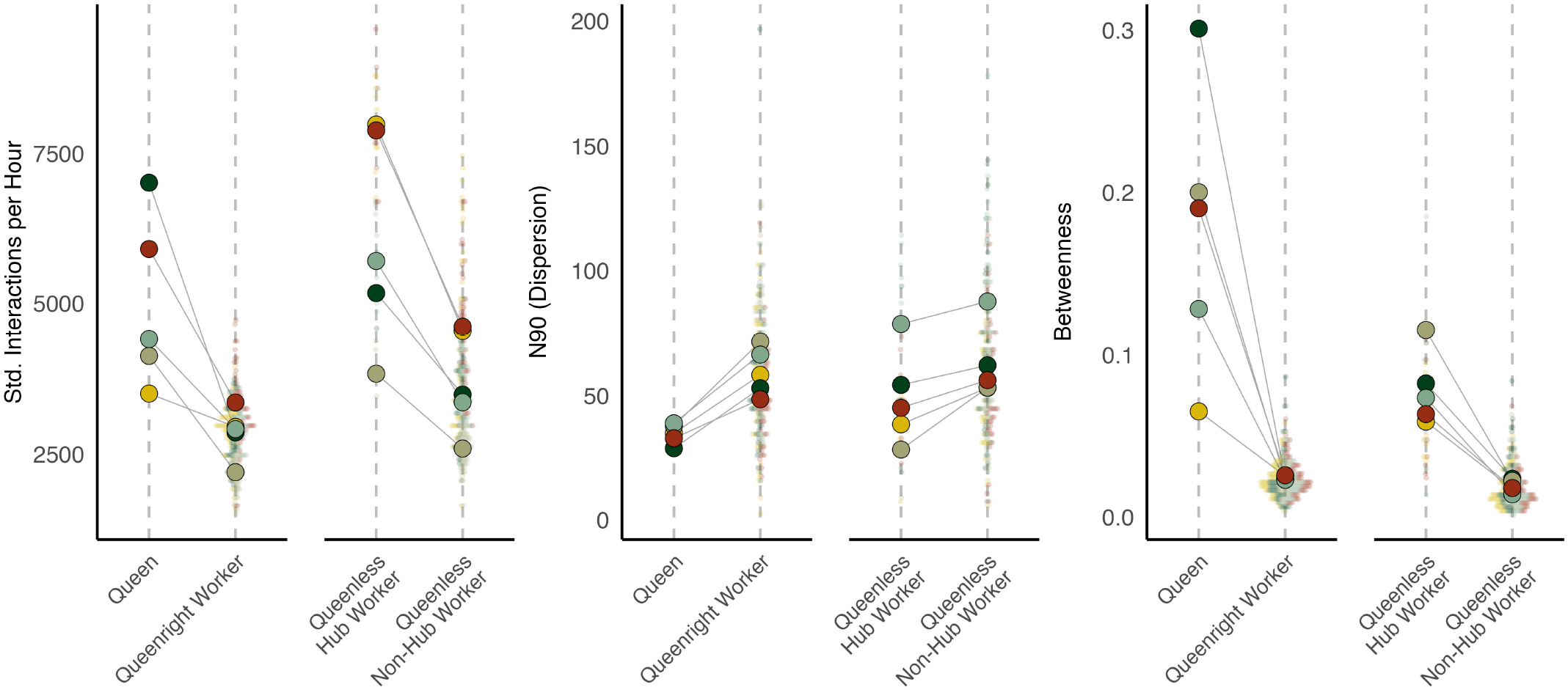
Key network parameters standardized interaction rate (a), dispersion (b), and betweenness centrality (c) in queens, queenright workers, queenless hub workers, and queenless non-hub workers. Large dots show colony means and smaller dots show individual measurements for the parameters.

**Figure S15.**
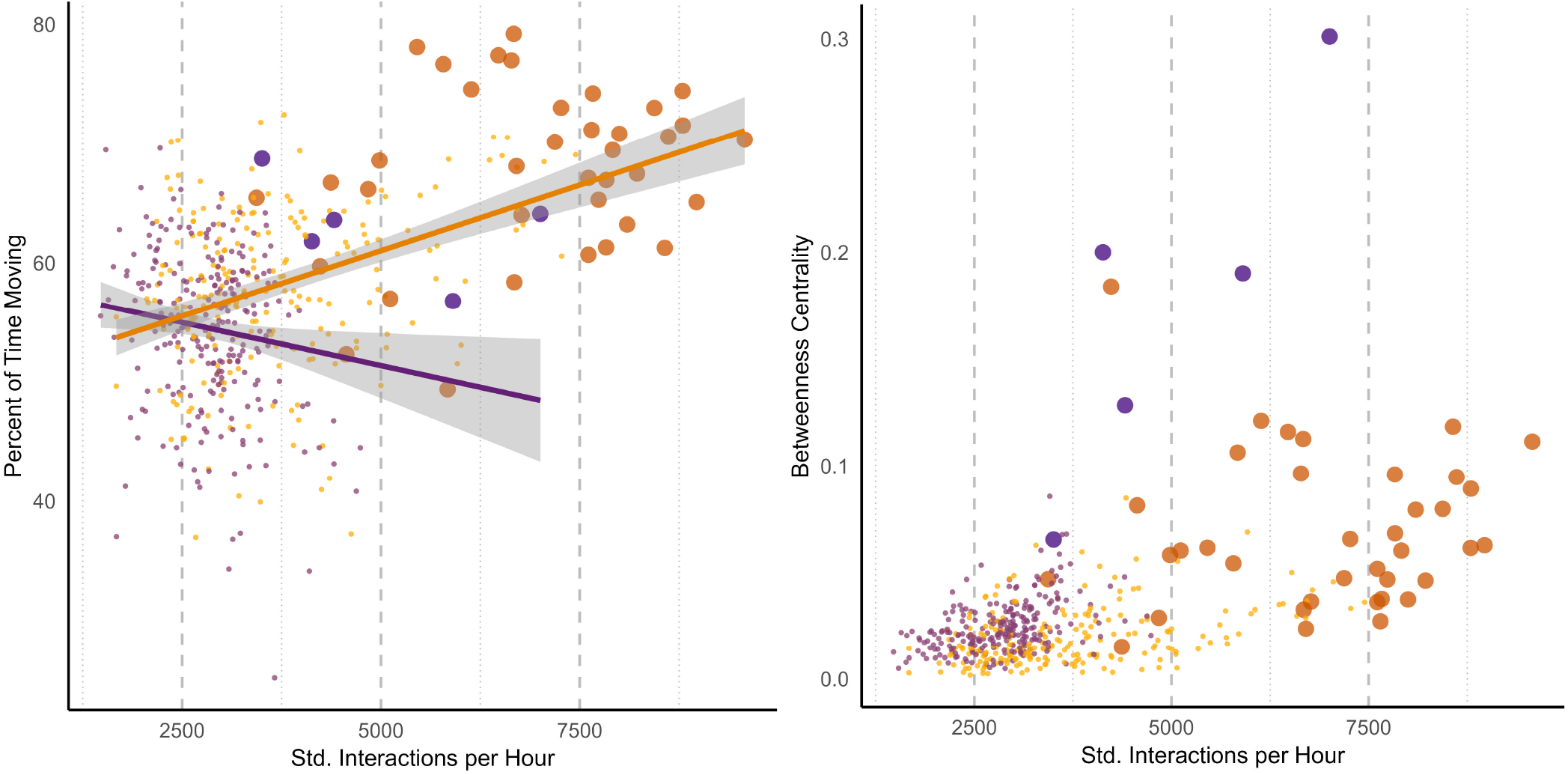
(left) Percent of time moving by standard interactions per hour and (right) betweenness centrality by standard interactions per hour. Purple corresponds to queenright individuals and orange to queenless, enlarged purple dots are queens and enlarged orange dots represent queenless hub bees.

**Figure S16.**
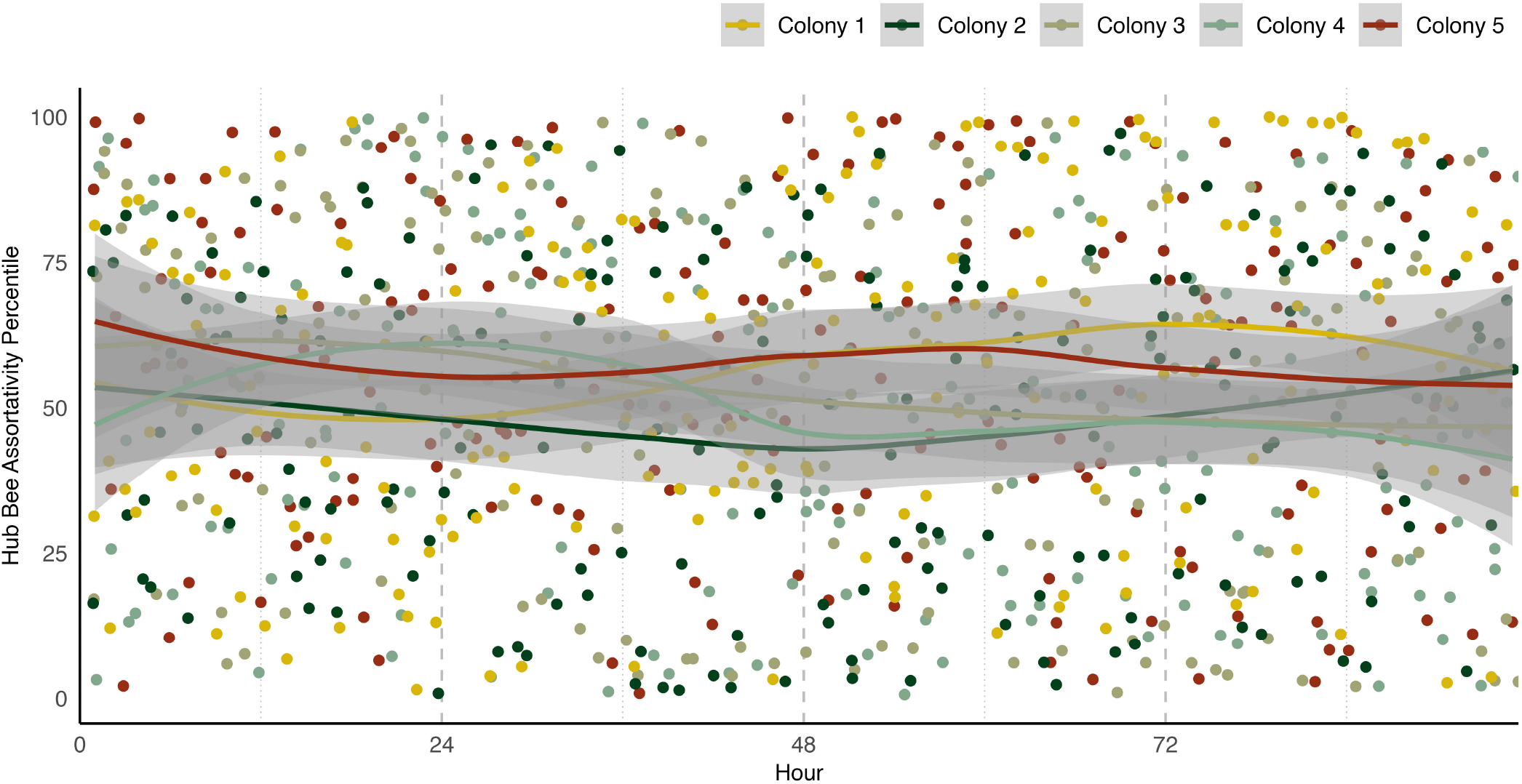
Assortativity between hub bees in queenless colonies over time. Colonies are split by color and LOESS fit for each colony is shown.

**Figure S17.**
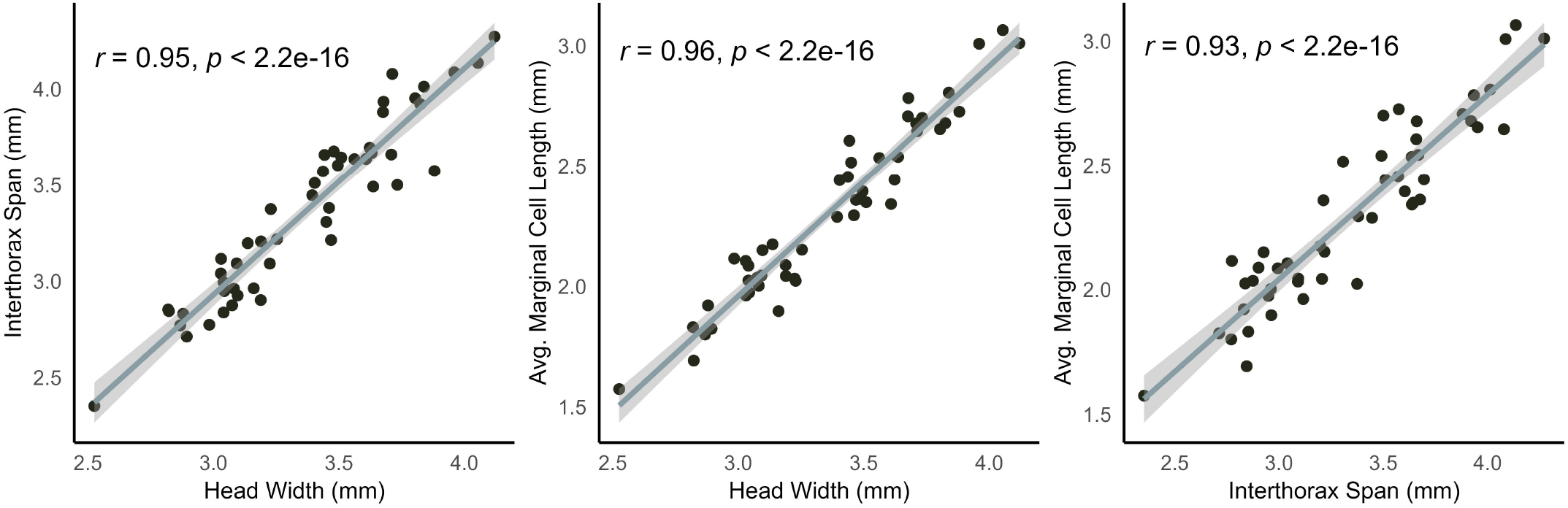
Scatter plots showing relationships between size measurements. (left) Scatter plot showing relationship between interthorax span (mm) and head width (mm). (center) Scatter plot showing relationship between average marginal cell length (mm) and head width (mm). (right) Scatter plot showing relationship between average marginal cell length (mm) and interthorax span (mm).

